# Grid-cell modules remain coordinated when neural activity is dissociated from external sensory cues

**DOI:** 10.1101/2021.08.29.458100

**Authors:** Torgeir Waaga, Haggai Agmon, Valentin A. Normand, Anne Nagelhus, Richard J. Gardner, May-Britt Moser, Edvard I. Moser, Yoram Burak

**Author notes:** Equal contributors (Torgeir Waaga, Haggai Agmon).

## Abstract

The representation of an animal’s position in the medial entorhinal cortex (MEC) is distributed across several modules of grid cells, each characterized by a distinct spatial scale. The population activity within each module is tightly coordinated and preserved across environments and behavioral states. Little is known, however, about the coordination of activity patterns across modules. We analyzed the joint activity patterns of hundreds of grid cells simultaneously recorded in animals that were foraging either in the light, when sensory cues could stabilize the representation, or in darkness, when such stabilization was disrupted. We found that the states of different grid modules are tightly coordinated, even in darkness, when the internal representation of position within the MEC deviates substantially from the true position of the animal. These findings suggest that internal brain mechanisms dynamically coordinate the representation of position in different modules, to ensure that grid cells jointly encode a coherent and smooth trajectory of the animal.

## INTRODUCTION

Recently, techniques that enable simultaneous recording of activity in dozens to hundreds of neurons (Ghosh et al., 2011; Jun et al., 2017; Steinmetz et al., 2021; Zong et al., 2017) have enabled a shift from the measurement of single cell activity in relationship to external correlates, to investigation of the joint population activity patterns in large neural ensembles. This change of perspective has led to various attempts to characterize neural activity patterns as residing within restricted, low dimensional spaces using linear (Gallego et al., 2018; Mazor and Laurent, 2005; Stringer et al., 2019) or non-linear (Chaudhuri et al., 2019; Gardner et al., 2021; Rubin et al., 2019; Rybakken et al., 2019) dimensionality reduction techniques. One of the most striking outcomes of these attempts has emerged in neural circuits involved in the representation of an animal’s position relative to the environment. In several such circuits in flies and mammals, neural activity patterns have been shown to robustly reside in low-dimensional nonlinear manifolds, even when the neural activity is dissociated from external inputs to the network (Chaudhuri et al., 2019; Gardner et al., 2021; Kim et al., 2017; Rybakken et al., 2019; Seelig and Jayaraman, 2015). This finding opens up the possibility to decode the low-dimensional variable that is represented within these circuits, and to examine how the brain utilizes such representations across multiple sub-circuits to implement computational functions.

Here we examine the dynamics of grid cells in the medial entorhinal cortex (MEC). Grid cells exhibit multiple firing fields as a function of an animal’s spatial location. The fields are arranged on a hexagonal lattice in open field environments (Hafting et al., 2005). Within each individual animal, grid cells are allocated to discrete modules, each defined by a common grid spacing and angular orientation (Barry et al., 2007; Stensola et al., 2012). Jointly, the activity of grid cells across multiple modules implements a highly efficient population code for position (Burak, 2014; Fiete et al., 2008; Mathis et al., 2012; Sreenivasan and Fiete, 2011; Welinder et al., 2008).

The spatial tuning curves of individual grid cells indicate that grid cell population activity within each module is confined to lie on a low-dimensional manifold with toroidal topology (Gardner et al., 2021). Accumulating evidence has suggested that this confinement is achieved through network mechanisms within the MEC, under diverse behavioral conditions and independently of inputs from other brain regions. Early evidence came from observing the correlation structure of activity in pairs of cells: phase relationships between grid cells within a module are tightly preserved over time and across environments (Fyhn et al., 2007; Yoon et al., 2013). The phase relationships are maintained also during sleep (Gardner et al., 2019; Trettel et al., 2019) and under hippocampal inactivation, despite the absence of a grid-like spatial response pattern (Almog et al., 2019). Very recently, simultaneous recordings of spiking activity in dozens of cells provided direct evidence that neural activity patterns are closely confined to two dimensional manifolds with toroidal topology, which are tightly preserved across environments and in sleep (Gardner et al., 2021). Thus, grid cells within a module encode together a two-dimensional quantity which, in some conditions, could be dissociated from the true position of the animal.

All of the above findings are in agreement with predictions made by continuous attractor network (CAN) theory (Burak and Fiete, 2009; Fuhs and Touretzky, 2006; Guanella et al., 2007; McNaughton et al., 2006). According to this theory, grid cells within each module are recurrently connected, and thus form a sub-network within the MEC. The recurrent synaptic connectivity within each module constrains the joint activity of cells to a restricted, but continuous repertoire of possible coactivation patterns which is stable across behavioral states and conditions, even in the absence of sensory inputs.

Single modules alone, however, cannot represent a unique position of an animal within a typical environment. It is necessary to consider the coordination of activity across modules in order to assess how grid cells encode the brain’s internal representation of position. The question of coordination becomes especially important under conditions in which sensory cues are poor or absent (Burak, 2014). Since population activity of an individual module lies on a two-dimensional manifold, the joint activity of *M* modules spans, at least in principle, a 2*M* dimensional space. However, during continuous motion in a given environment, and in the presence of salient sensory cues, the state of each module is faithfully mapped to the location of the animal in two-dimensional space. Hence, under continuous motion of the animal, the joint population activity patterns of multiple modules span a highly restricted two-dimensional subspace of the full 2*M* dimensional space. This raises the question, whether the states of different modules are updated in a similarly coordinated manner when the states of individual modules are dissociated from the true position of the animal, e.g., in the absence of salient sensory cues.

The coordination of activity across grid cell modules is highly consequential from the perspectives of neural coding and dynamics. The modular structure of the grid cell code for position confers it with large representational capacity (Burak, 2014; Fiete et al., 2008; Mosheiff and Burak, 2019; Sreenivasan and Fiete, 2011; Welinder et al., 2008). Yet, within a given environment, the modularity of the grid cell code poses a significant challenge for the neural circuitry that maintains the representation and updates it based on self-motion. Under conditions in which sensory inputs are absent or poor, the representation of position in individual grid cell modules might drift relative to the actual position of the animal. If these drifts are not identical in different modules, they would rapidly lead to combinations of spatial phases that do not represent any position in the vicinity of the animal, resulting in abrupt shifts in the represented position. Thus, independent drifts lead to catastrophic errors when activities are read out from multiple grid cell modules, and would therefore be highly detrimental for the coding of position by grid cell activity. The difficulty arising from occurrence of such catastrophic readout errors has been identified in early works on grid cell coding (Fiete et al., 2008). Since then, two solutions have been proposed. In one solution (Agmon and Burak, 2020; Sreenivasan and Fiete, 2011; Welinder et al., 2008) the hippocampal network reads out the position represented by grid cells, and feedback projections from hippocampus to the MEC correct small incompatible drifts accrued in each of the modules. A second solution (Kang and Balasubramanian, 2019; Mosheiff and Burak, 2019) involves synaptic connectivity between modules.

Empirically however, very little is known about the relationship between population activity patterns of grid cells across distinct modules. Previous research has focused on coactivation patterns within modules for two reasons: first, simultaneously recorded cells using tetrodes often belonged to the same module. Second, coactivation patterns of inter- and intramodule grid cell pairs are fundamentally different. Grid cells within a module maintain strict relationships in their activities that can be probed by analyzing the joint activity in pairs of simultaneously recorded cells. On the other hand, the activity of two cells that belong to different modules might be correlated or anti-correlated depending on the animal’s position, even within a fairly small environment. Due to this lack of an expected rigid correlation (or anti-correlation), it is difficult to characterize inter-module coordination based on pair recording analysis. In order to identify higher order dependencies across modules, it is necessary to decode activity from multiple cells within each module – requiring larger numbers of simultaneously recorded cells from multiple modules, which have only recently become available.

Here, using Neuropixels silicon probes (Jun et al., 2017; Steinmetz et al., 2021) we recorded the simultaneous activity of grid cells from multiple modules with dozens of units in each module. Rats were deprived of visual cues in order to test whether the internal representations of position in distinct modules remain coordinated even when dissociated from the animal’s true position. By decoding the simultaneous grid cell activity, we demonstrated that grid cell modules retain, to a high extent, coordination even when the mapping between grid cell activity and position deteriorates. These results indicate that network mechanisms within the brain coordinate the activity of different modules, independently of external sensory inputs.

## RESULTS

We recorded spiking activity from rats foraging in a circular arena with a diameter of 150 cm under light and complete dark conditions (Fig. 1a). The arena was cue-less except for a single vertical cue card at a fixed location along its circumference which was visible in light and completely invisible in darkness, and tactilely inaccessible. The circular arena was rotationally symmetric, thus minimizing the information about absolute position coming from encounters with the walls (Hardcastle et al., 2015; Keinath et al., 2018). The arena was surrounded by a floor-to-ceiling blockout blind to eliminate access to distal visual cues, and the experimental protocol was designed to minimize other positional cues (*Online Methods*).

**Fig. 1:**
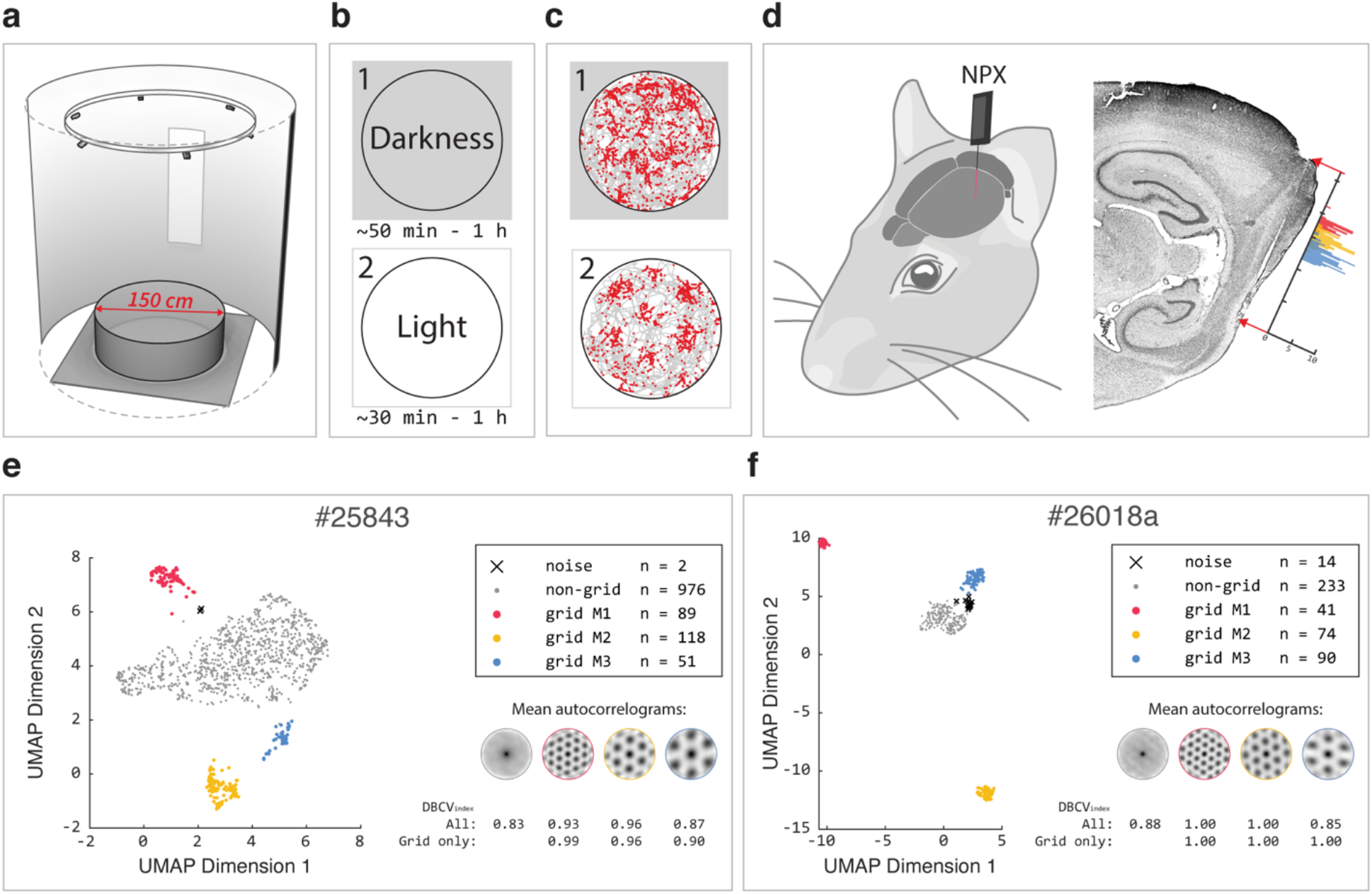
Experimental setup and module classification. **a,** Recording arena: electrophysiological recordings of the spiking activity in the MEC were collected while the rat ran freely in a 150 cm diameter cylindrical arena surrounded by a floor-to-ceiling blind and a single fixed and tactile inaccessible cue card. **b,** Protocol: first the animals ran in darkness, then in the same arena with the lights on. **c,** The spikes of a representative grid cell from the most dorsal module (red distribution in (d)) are superimposed in red on the traveled path of the rat in gray, in the darkness (1) and the light task (2). **d,** Left: illustration of implantation site for neuropixels probe. Right: sagittal section of the rat brain (#26018), showing the probe shank through the superficial layers of MEC. The histogram shows the grid cell count across dorso-ventral recording depths from three modules. The distance between two adjacent ticks along the probe shank axis corresponds to 1 mm. **e-f,** Module classification for the two recording sessions with the largest number of simultaneously recorded grid cells. Left: for each recording, scatterplots show the two-dimensional UMAP (McInnes et al., 2018) projection of all recorded units’ autocorrelograms. Each point is color coded by it’s DBSCAN cluster assignment. Right: mean autocorrelograms for each cluster and validity (DBCV) index (*Online Methods*) including or excluding the non-grid cluster.

Data was collected from four animals and an overall of five recording sessions, each consisting of a 50-60 minutes recording in the dark immediately followed by a 30-60 minutes recording in the light (Fig. 1b-c). Neuropixels probes were implanted in MEC (Fig. 1d and Supplementary Fig. 1). Out of 3310 recorded cells with >500 spikes in the 5 light trials, 842 grid cells were identified and classified into modules as described in *Online Methods* and in (Gardner et al., 2021). Briefly, a non-linear dimensionally reduction technique (McInnes et al., 2018) was applied to feature vectors derived from the spatial autocorrelation of each cell rate map, followed by clustering (Ester et al., 1996). The procedure ensures modular separation as grid cells with similar auto-correlograms, and thus similar spacing and orientation, are separated into distinct clusters while cells lacking a spatially periodic tuning feature, are separated from the grid cell clusters. In addition to overcoming problems from traditional classification methods caused by skewed grid patterns, false positives would only occur in the unlikely event that the autocorrelogram of a non-grid cell randomly has a pattern with the same spacing and orientation as the real grid cells. Using this procedure, a clear and unambiguous clustering of grid cells into modules was observed in all the sessions (Fig. 1e-f and Supplementary Fig. 2). In four out of five recording sessions, many simultaneously recorded grid cells (ranging from 31 to 118 in individual modules) were obtained from three distinct modules, and in one session such data was obtained from two distinct modules (Table 1).

**Table 1:** Numbers of simultaneously recorded grid cells (allocated to modules) for each recording session.

| Recording session | Module 1 | Module 2 | Module 3 |
| --- | --- | --- | --- |
| #25843 | 89 | 118 | 51 |
| #26018a | 41 | 74 | 90 |
| #26018b | 40 | 68 | 78 |
| #26820 | 31 | 44 | 35 |
| #26718 | 47 | 36 | - |

Several measures indicated that the association between grid cell activity and the position of the animal deteriorated in the sensory deprived condition. The characteristic periodicity of grid cells was significantly disrupted in the dark trials (Fig. 2a and Supplementary Fig. 3). Gridness score and information content were considerably reduced compared to the baseline light trials (Fig. 2b-c). In addition, we decoded position from the population activity patterns, and compared the magnitude of decoding errors in the light and dark trials. We have done so using two types of decoders that are used later in the manuscript, and are described in *Online Methods*. The Mean Absolute Error (MAE) of the decoded position relative to the animal’s true position was substantially larger in dark trials than in baseline light trials (Fig. 2d-e). This was consistent when the decoding of simultaneous spike trains from dark trials was performed either using light- or dark-generated rate maps (Supplementary Fig. 4), and across the two decoders.

**Fig. 2:**
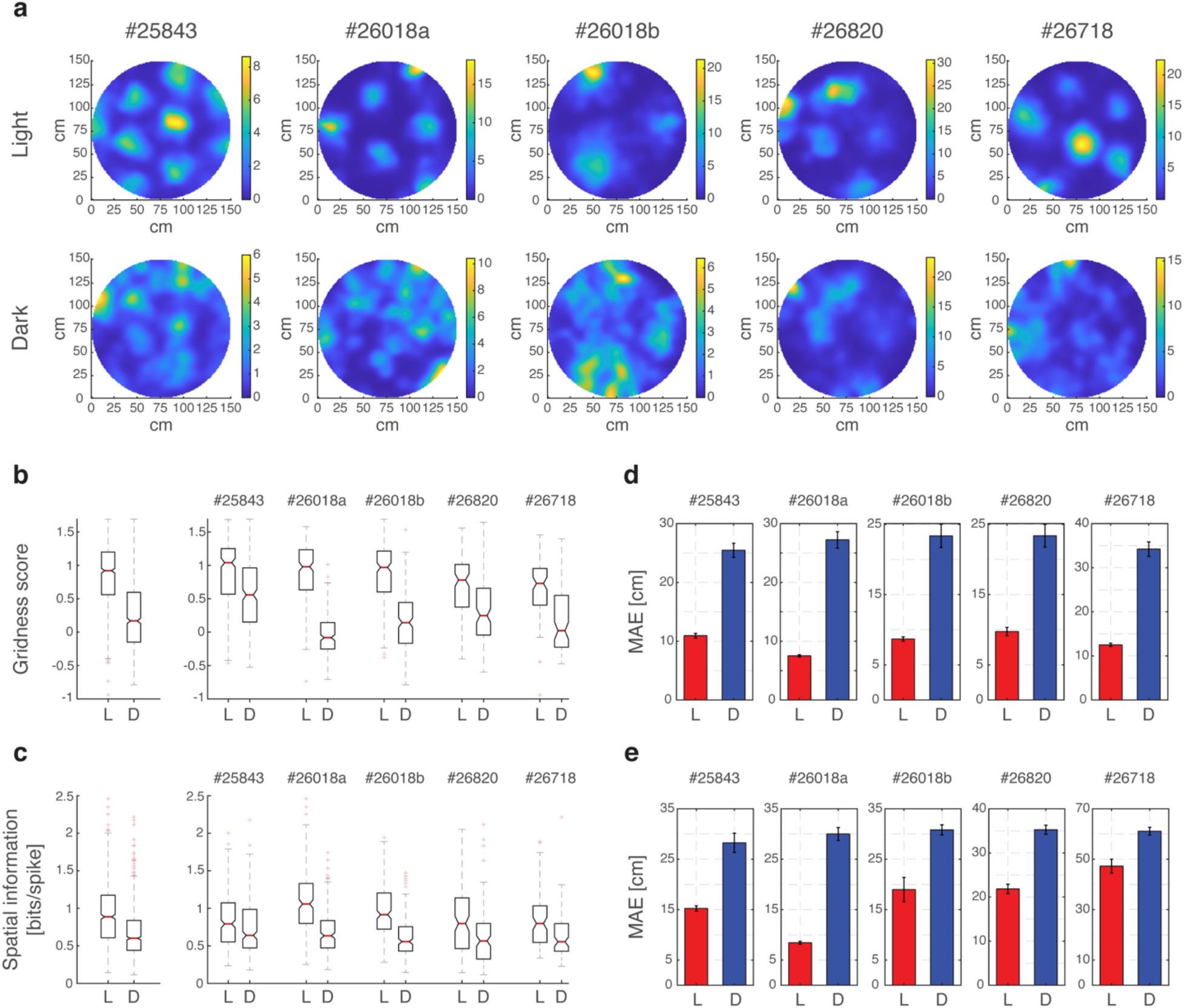
Grid cell spatial responses deteriorate in darkness. **a,** Example of rate maps from each session for light and darkness conditions. Three more examples from each recording session are shown in Supplementary Fig. 2. **b,** Gridness scores in light and darkness for all cells across all recording sessions (left), and for all cells from single recording sessions (right). **c,** Spatial information in light and darkness for all cells across all recording sessions (left), and for all cells from single recording sessions (right). **d,** Mean Absolute Error (MAE) of the Markov decoder (*Online Methods)* in light (red) and darkness (blue) for each recording session. Error bars are ±SEM. **e,** Same as (d) but for the kernel decoder (*Online Methods*).

### Pairwise correlations

As a first step to address the question of inter-module coordination, we considered pairwise correlations of the spiking activity, smoothed with a 50 ms Gaussian kernel (*Online Methods*), in similarity to previous works that were based on tetrode recordings (Almog et al., 2019; Chen et al., 2016; Fyhn et al., 2007; Gardner et al., 2019; Pérez-Escobar et al., 2016; Trettel et al., 2019; Yoon et al., 2013). Differences in the correlation structure of inter-versus intra-module pairs were expected under light conditions for the following reason: the activity of intra-module grid cell pairs is either correlated or anti-correlated irrespective of the position of the animal, whereas the activity of inter-module grid cell pairs can be correlated in some parts of the environment and uncorrelated in others (Fig. 3a). Consequently, weaker absolute spiking correlations were expected on average in inter-module pairs compared to intra-module pairs. Indeed, under light conditions intra-module pairs showed higher absolute spiking correlations compared to intermodule pairs (Fig. 3b). The observed zero-lag correlations could be explained quite well by the cells’ rate maps and the animal’s trajectory, even in inter-module pairs (Fig. 3c). Thus, the correlations observed in the spiking activity of inter-module pairs were weak but still driven, to a large extent, by their spatial selectivity.

**Fig. 3:**
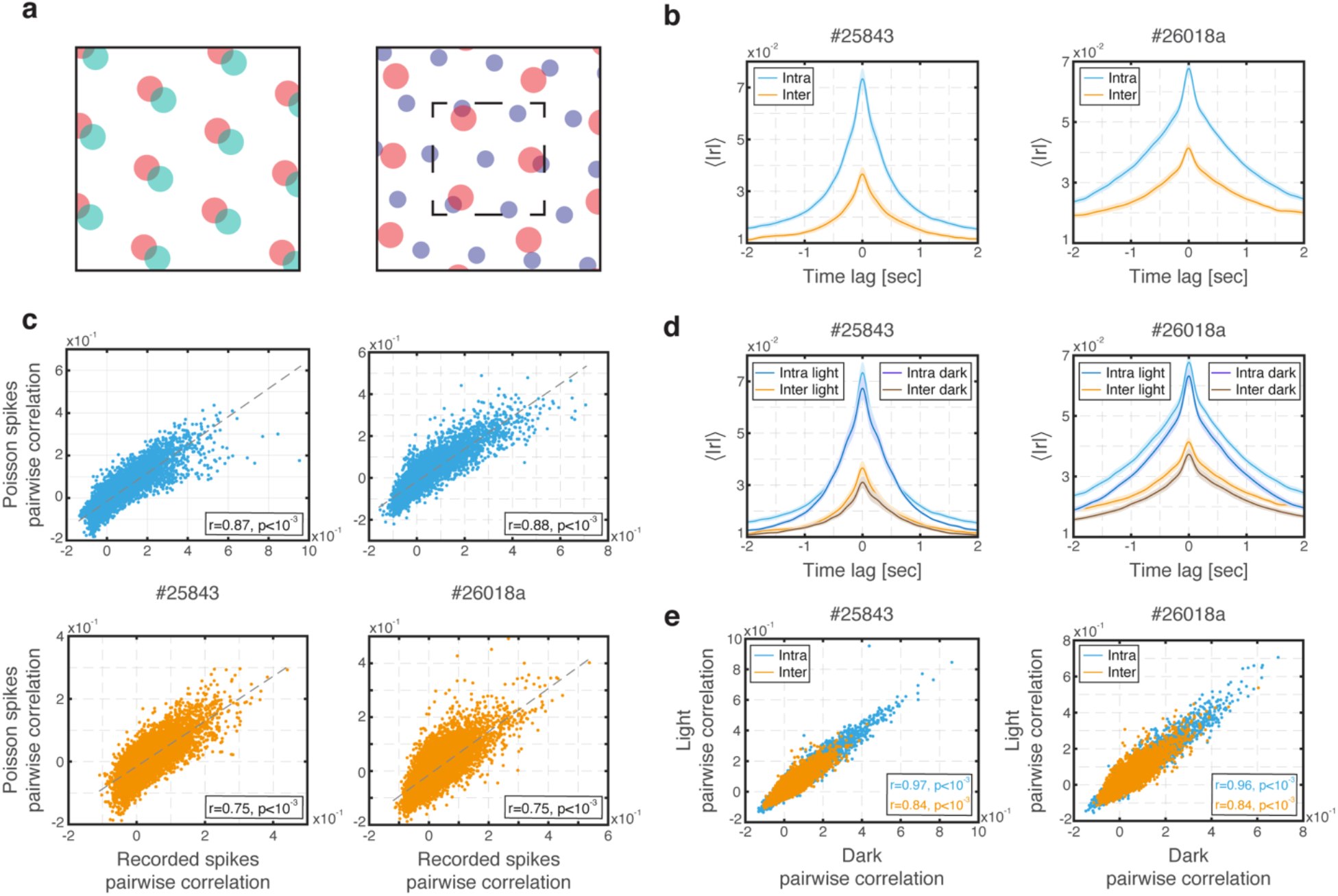
Spiking correlations of intra- and inter-module pairs. **a,** Schematic illustration showing spatial tuning curves of two intra-module (left, red and turquoise) and two inter-module (right, blue and red) grid cells. Firing fields of intra-module pairs (left) either overlap throughout the whole environment (as shown in the figure) causing temporally correlated firing or are disjoint throughout the whole environment causing anti-correlated firing. On the other hand, in intermodule pairs (right) the degree of correlation (or anti-correlation) between firing fields varies in different parts of the environment: firing fields overlap inside the small dashed square, and are disjoint elsewhere. Consequently, absolute spiking correlations tends to be weaker in intermodule pairs than in intra-module pairs. This effect is more pronounced in large environments compared to small environments (Supplementary Fig. 5). **b,** Absolute cross-correlation (Pearson coefficient) of inter- and intra-module light spiking activities, averaged over cell pairs, in the two recording sessions with the largest number of simultaneously recorded neurons (left and right panels). Shaded error bars are ±SEM. **c,** Pairwise correlations of all possible intra- (top) and inter- (bottom) module pairs from recorded light spiking activity versus Poisson generated spikes using measured rate maps and the corresponding recorded light trajectory. Correlation coefficient and p-value are specified in the inset. **d,** Same as (b) but with superimposed inter- and intra-module spiking activities from darkness. **e,** Pairwise correlations of all possible inter- and intra-module pairs from recorded light spiking activity versus recorded dark spiking activity.

Having established that spiking correlations are related to spatial selectivity both in intra- and inter-module pairs, we next compared pairwise spiking correlations in dark and light trials. We reasoned that if, in darkness, grid cells cease to consistently encode a unique spatial location, their spiking correlations would diminish. The absolute magnitude of the spiking correlations in *intro*-module pairs was similar, on average, to those observed in the light (Fig. 3d) even when, in the dark, the spatial stability of single grid cell representations was disrupted. This is in accordance with recordings performed in mice in dark environments (Chen et al., 2016; Pérez-Escobar et al., 2016) and with results found in sleep (Gardner et al., 2019), and as expected based on CAN models. However, the average absolute magnitude of spiking correlations in *inter* module pairs was similar in light and dark conditions as well. Furthermore, zero-lag correlations were preserved between light and dark conditions at the level of individual cell pairs both for inter- and intra-module pairs (Fig. 3e).

The preservation of inter-module spiking correlations in the dark supports the hypothesis that modules maintain coordination even in the absence of sensory cues. However, it is difficult to interpret this result quantitatively, since inter-module spiking correlations are already low even in the light, and may be influenced also from sources other than the correlation between their spatial receptive fields, such as co-fluctuation of firing rates in the entire population (Okun et al., 2015). Analysis of our simultaneous recordings from dozens of grid cells per module (Table 1) could potentially overcome these limitations, by revealing higher order dependencies in the activity of cells from different modules that are not strongly evident in spiking correlations within pairs of cells. Therefore, we next analyzed the recorded simultaneous population activities, using two complementary approaches.

### Likelihood of the simultaneous population spike trains

In the absence of sensory cues and under the hypothesis of inter-module coordination, a unique position should be coherently represented by the joint activity of grid cells in different modules, even when spatial firing patterns of individual grid cells seem disrupted. When the joint representation is read out under these conditions, it can, however, deviate substantially from the animal’s true position compared with baseline light trials. We therefore sought to identify a measure for the coherence of the joint simultaneous spike trains, which is *independent* of the animal’s true position.

In the first analysis approach, we derived the likelihood of the simultaneously recorded spike trains, summed over all possible trajectories (thus, independently of the actual trajectory) under simple assumptions that are outlined below. This likelihood is written as

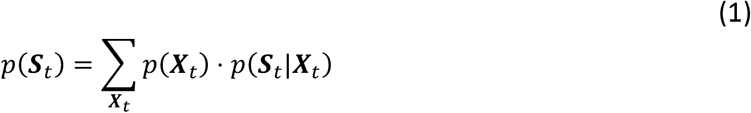

where ***S**_t_* represents the simultaneous spike trains emitted by all the neurons in the population from the beginning of the experiment up to time *t*, and ***X**_t_* represents a particular trajectory of the animal. The likelihood of the spike trains conditioned on the trajectory, *p*(***S**_t_|**X**_t_*), is evaluated under the assumption of Poisson firing, with a rate which is determined by ***X**_t_* and by the tuning curves of all the neurons in the population. Finally, we assumed that the trajectories are continuous (following random walk statistics for simplicity), enforced through the prior *p*(***X**_t_*) (see also *Online Methods*).

On the right hand side of Equation (1), a probability is assigned to each particular realizable trajectory irrespective of the spike trains, followed by a multiplication with the probability of the simultaneously recorded spike trains conditioned on this particular trajectory. This probability is subsequently averaged over all possible trajectories, weighted by the prior. The outcome *p*(***S**_t_*) can be interpreted as a measure which describes the likelihood that the simultaneously recorded spike trains represent some continuous (yet unknown) trajectory, drawn from the prior distribution. Importantly, in a practical implementation there is no need to explicitly calculate the specific probabilities for each of the possible trajectories, which would be unfeasible. Instead, we derived a simpler, exact analytical expression for the average log likelihood per time bin, denoted by *L* (*Online Methods*, Eq. 3). The evaluation of *L* using this expression relies on the Markov property of the spiking model and the prior *p*(***X**_t_*), and involves decoding of the spiking activity using a *Markov decoder* (*Online Methods* and *Supplementary Information*).

In the expression for the likelihood (Eq. 1), it is assumed that all the neurons fire in response to the same trajectory ***X**_t_*, regardless of the identity of the module to which they belong. If this assumption is correct then this trajectory, as well as nearby trajectories, will make a large contribution to the likelihood (Fig. 4a). However, if different modules accrue drifts independently in the dark and thus represent internally different trajectories, there will be no single trajectory in the sum within Eq. 1 that makes a large contribution to the likelihood, and we expect the sum to be significantly smaller. Thus, the likelihood introduced above (Eq. 1) can serve as a measure of coherence.

**Fig. 4:**
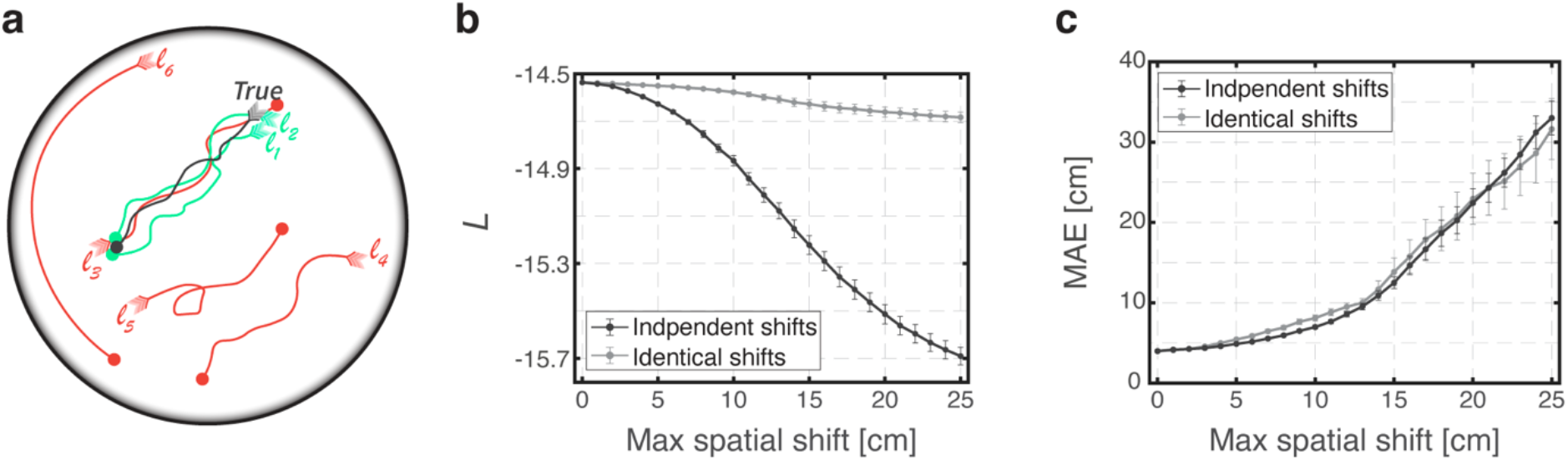
Likelihood of simultaneous spike trains can serve as a measure of coordination across modules. **a,** Schematic illustration demonstrating the likelihood approach with few illustrated trajectories, each starting from an arrowhead and ending in a point. The true trajectory of the animal is the black trace. The likelihood of the simultaneous spike trains is evaluated independently for each specific trajectory, and is averaged across all possible trajectories. Few possible representative trajectories are illustrated: trajectories *l*_1_ and *l*_2_ (turquoise) are two trajectories which are close to the true trajectory and thus have a high likelihood based on the spiking activity. Trajectory *l*_3_ is very similar to trajectories *l*_1_ and *l*_2_ but has low likelihood (red) since it goes in the opposite direction, thus having reversed temporal structure. Trajectory *l*_4_ is an identical copy of trajectory *l*_2_ but at a different part of the arena thus also having low likelihood (red). Trajectories *l*_5_ and *l*_6_ are two other trajectories with low likelihood (red). **b,** Likelihood of simulated Poisson spikes using measured rate maps and the recorded light trajectory from recording session #26018b, evaluated versus varying magnitudes of spatial shifts. When shifts are applied independently in each module, the likelihood decreases significantly but it decreases only slightly (due to boundary conditions) when these shifts are identical. Error bars are ±SEM. **c,** The corresponding Mean Absolute Error (MAE) of the decoder. The MAE increases significantly both for independent and for identical spatial shifts as their magnitude increases. Error bars are ±SEM.

To validate that the likelihood can be used to distinguish between the scenarios of coordinated and uncoordinated drifts across modules, we first analyzed simulated spike trains. All neurons in the simulated data fired according to the same recorded trajectory, and based on measured tuning curves which were taken from one of our datasets. Therefore, the different modules were precisely coordinated in the simulation. To mimic the consequences of uncoordinated drifts across modules in a way that can be applied also to recorded datasets, we introduced during the decoding process artificial spatial shifts in the neuron’s rate maps (*Online Methods*) that, for simplicity, were constant throughout each simulation. Such shifts were identical within each module but drawn randomly and independently in different modules (and were thus uncoordinated across modules).

We observed that applying independent spatial shifts reduced the likelihood (Fig. 4b, black trace). As expected, such shifts also increased the MAE of the decoder with respect to the true position (Fig. 4c, black trace). On the other hand, the likelihood was nearly unaffected when the artificial spatial shifts were identical across all modules, even though the MAE with respect to the true position increased significantly (Fig. 4b,c gray traces). The small reduction in the likelihood seen in Figure 4b (gray trace) is due to boundary effects, and vanishes when such effects are eliminated (Supplementary Fig. 6). The results shown in Fig. 4 confirmed that the average log likelihood could be used as a measure of coherence of the simultaneous spike trains.

We next applied the likelihood-based approach to recorded datasets from the light and dark conditions (5 recordings sessions from 4 animals; Table 1), in order to assess whether modules remain coordinated in the dark. If, in the dark, the phases of individual modules accrued independent drifts relative to the animal’s true position, a reduction in the likelihood would be expected relative to the light. In order to faithfully compare likelihoods between light and dark conditions, it was necessary to take into account modifications in the mean firing rates of individual neurons across the two conditions: specifically, mean firing rates were more likely to reduce than increase in darkness, resembling previous results from mice (Pérez-Escobar et al., 2016). Thus, we evaluated *rote-odjusted* likelihoods of dark and light simultaneous spike trains, obtained by down-sampling spikes to match the mean firing rates between the two trials (*Online Methods* and *Supplementory Informotion*), henceforth referred for convenience as likelihood.

We found that the likelihood of simultaneously recorded spike trains in light trials was slightly higher than the likelihood in dark trials, even though the MAE was much larger in darkness than in baseline light trials (Fig. 5a, zero spatial shift). To demonstrate that the observed similarity in likelihoods was not simply an outcome of the rate-adjustment procedure, we also considered permuted rate-adjusted spike trains, which preserved the mean firing rates. Under such a permutation, the evaluated likelihood decreased drastically and the MAE drastically increased (Supplementary Fig. 7), demonstrating the importance of temporal structure of the simultaneous spike trains. To assess the significance of the small observed likelihood differences in relation to the question of module coordination, we evaluated the expected reduction in the likelihood under independent spatial shifts of varying magnitudes, as in Fig. 4 b-c. We found that the spatial shifts required in order to reduce the likelihood in the light to the value observed in the dark (point p1 in Fig. 5a left) would only generate a small increase in the MAE — of a few centimeters, whereas much larger spatial shifts would be required in order to match the actual MAE observed in the dark when using zero spatial shifts (point p2 in Fig. 5a right). Thus, the small reduction in the likelihood in the dark recordings at zero spatial shifts relative to the light is consistent with only small independent shifts across modules, of a few centimeters.

**Fig. 5:**
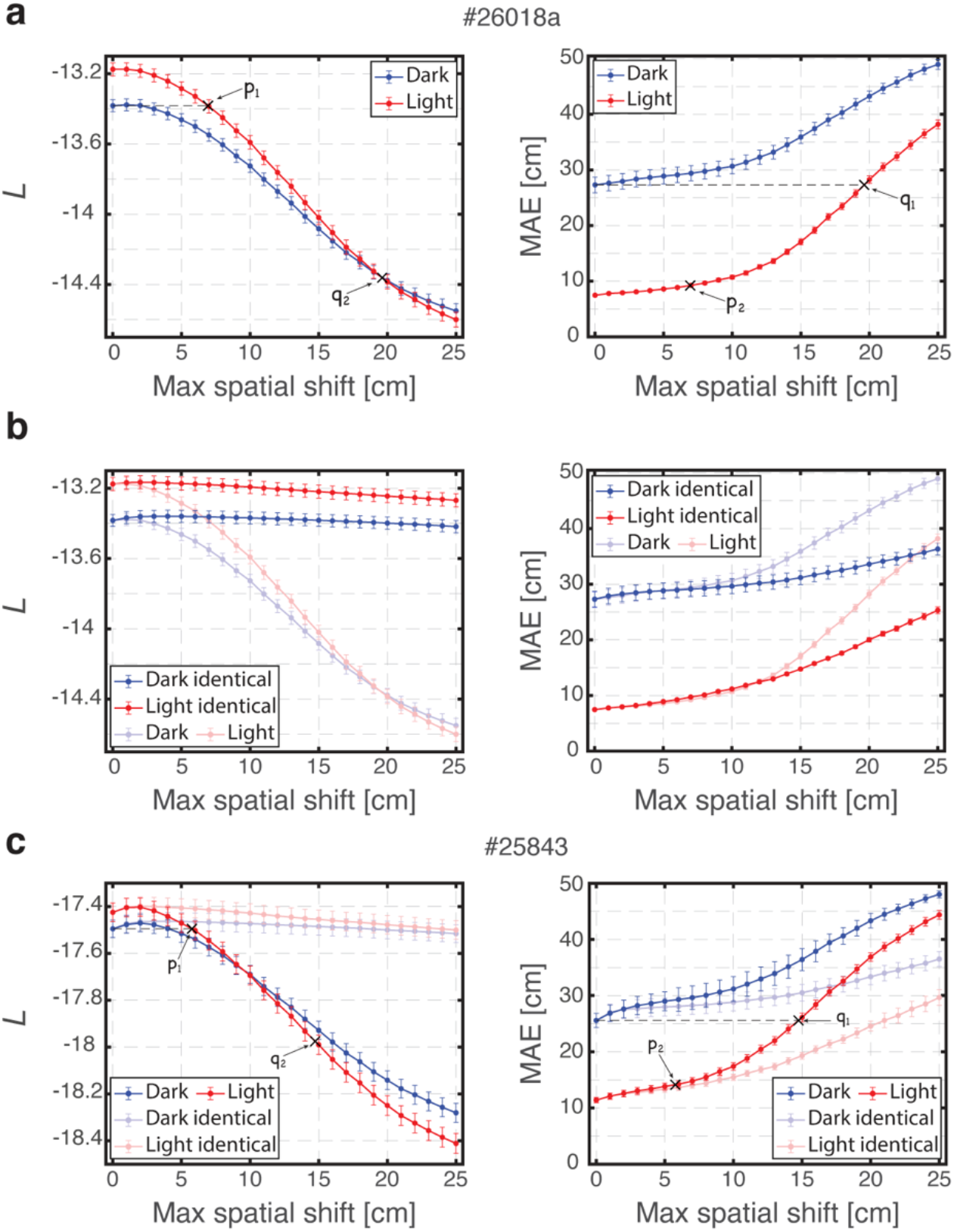
Analysis of likelihood of recorded simultaneous spike trains indicate coordination across modules. **a,** Likelihood of simultaneously recorded spike trains (left) and corresponding Mean Absolute Error (MAE, right) from dark and light trials, shown for varying magnitudes of independent module-wise spatial shifts applied on a single recording session (#26018a). Applying a maximal spatial shift of 6.9 cm in the light recording achieves the same likelihood as that of the dark recording with zero spatial shift (point p1, left panel), but generates only a slight increase of less than 2 cm in the corresponding MAE of the light recording relative to its zero spatial shift value (point p2, right panel). Conversely, applying a maximal spatial shift of 19.6 cm in the light recording achieves the same MAE as that of the dark recording with zero spatial shift (point q1, right panel), but generates a dramatic decrease in the likelihood (point q2, left panel). The difference in the likelihood between point q2 and the zero spatial shift point of the dark recording is much larger than the difference between the likelihood values of light and dark zero spatial shift points. Error bars are ±SEM. **b,** Same as (a) but with identical spatial shifts in all modules (full color traces; results for independent spatial shifts, same as in (a), are superimposed using faded colors for comparison). Under identical spatial shifts the dark and light likelihoods decrease only slightly (due to boundary conditions, left) while the corresponding MAE increases significantly in both cases (right). **c,** Same as (a) but for a another recording session (#25843), and with identical spatial shifts plotted in faded colors.

Conversely, the spatial shifts required in order to increase the MAE in the light to the value observed in the dark (point q1 in Fig. 5a right) would produce a substantial decrease in the likelihood if applied in an uncoordinated manner to the spike trains from the light recordings (point q2 in Fig. 5a left), to a value which is much smaller than the zero-shift likelihood observed in the dark recordings. Thus, the increase in the zero-shift MAE observed in the dark recordings must arise mostly from coordinated drifts across the modules. To validate that coordinated drifts across modules can increase the MAE without significantly reducing the likelihood, we introduced identical spatial shifts across all modules as in Fig. 4b-c. As expected, the likelihoods of dark and light trials were only slightly reduced while the corresponding MAE’s increased substantially (Fig. 5b). As in Fig. 4b, the small reduction observed in the likelihoods with the introduction of coordinated spatial shifts is due to boundary conditions.

Similar results from additional datasets are shown in Fig. 5c and in Supplementary Fig. 8.

### Decoding from individual modules

The likelihood-based approach described above can be applied to datasets with moderate numbers of cells per module, where the decoded position is very noisy. In most of our datasets the large number of simultaneously recorded cells enabled testing for inter-module coordination using a more direct approach, based on the decoding of position from individual modules.

In this approach, schematically illustrated in Fig. 6a, an internal multi-module representation of position, denoted by 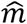, was first estimated by decoding spiking activity from all the grid cells. We used a *kernel decoder*, which estimates position based on spikes within a fixed time window, unlike the Markov decoder which has access to the entire spiking history (*Online Methods*). As expected, the multi-module posterior typically exhibited a single prominent peak within the enclosure, which could potentially deviate from the true position of the animal (*X* in Fig. 6a). Next, spiking activity from each module was decoded separately. As expected, single module posteriors typically exhibited approximately periodic peaks. To resolve this ambiguity, the position which maximized the posterior within the local vicinity of the position 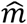 was selected as the corresponding decoded position 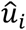, for each module *i* (*Online Methods*). Finally, the distances between the position 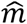 to each position 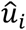 (denoted by *δ_i_*), and between 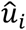 pairs (denoted by Δ_*ij*_) were evaluated and compared between dark and baseline light trials (Fig. 6a right).

**Fig. 6:**
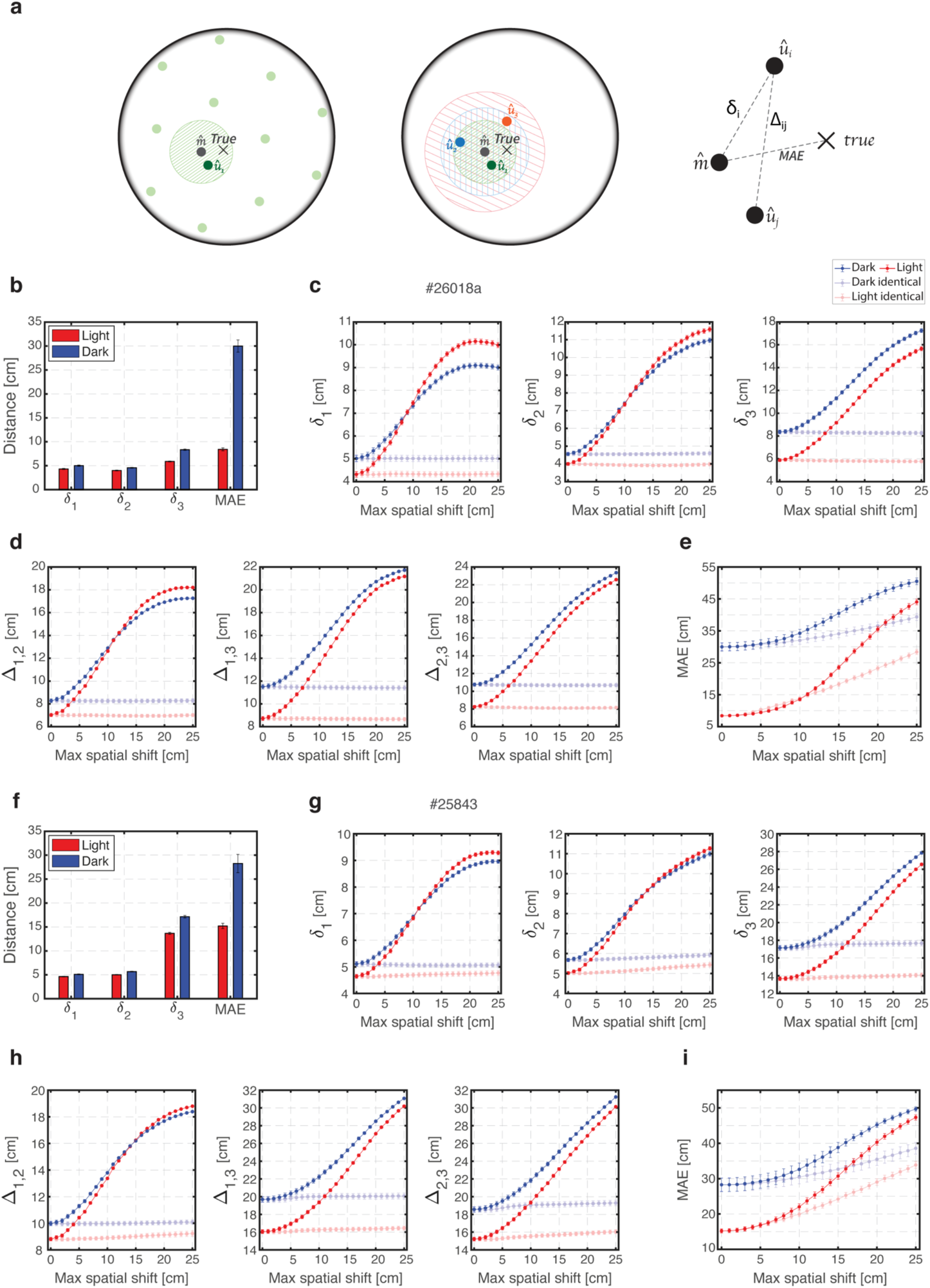
Decoding of population activity from individual modules reveals tight coordination of phases across modules in darkness. **a,** Schematic illustration of the uni-module decoding approach. Left: decoding the spiking activity from all grid cells typically produces a posterior with a unique blob in the arena, and the position of its maximum is chosen as the estimate of the multi module internal representation (gray circle 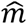). This estimation can deviate from the true position of the animal (black X symbol). Decoding the spiking activity from a single module typically produces a periodic posterior (faded green circles). The blob which is nearest to position 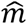 is selected (dark green circle), and the position of its maximum is chosen as the estimate of the uni-module internal representation, denoted by 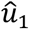. Middle: this procedure is repeated for each individual module (green, orange and blue) producing an estimated uni-module position for each module (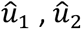 and 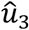). Right: the distance from position 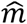 to the true position of the animal is averaged over time to produce the Mean Absolute Error (MAE). The mean distance between position 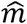 to each position 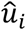 is defined as *δ_i_*, and the mean distance between each pair of positions 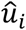 and 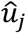 is defined as Δ_*ij*_. **b,** The measured MAE and distances *δ_i_* for dark and light from a single recording session (#26018a). The distances *δ_i_* in darkness are only ~1 cm larger than those in baseline light trials. Error bars are ±SEM. **c,** The distances *δ_i_* for varying magnitudes of module-wise independent spatial shifts and for identical spatial shifts. Increasing the magnitude of module-wise independent spatial shifts increases *δ_i_* dramatically, indicating that these measured distances *δ_i_* could potentially be much higher. The same magnitude of identical spatial shifts has no effect. Error bars are ±SEM. **d,** Same as (c) but for the corresponding distances Δ_*ij*_. **e,** Similar as (c) but for the corresponding MAE. The MAE increases for both module-wise independent and identical spatial shifts. **f-i,** Same as b-e but for another recording session (#25843).

The MAE of position 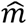, with respect to the animal’s true position, was higher in darkness than in baseline light trials (one example shown in Fig. 6b), indicating that the internal representation of position in the dark drifted relative to the true position. The distances *δ_i_* and Δ_*ij*_ were noisy and fluctuated in time both in dark and light conditions, but their mean was only slightly higher in darkness than in light (Fig. 6b). This is consistent with the results from the previous approach (Fig. 5), which demonstrated slightly higher likelihoods for simultaneously recorded light spike trains compared to corresponding dark trials.

Since the single module readout positions 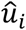 were restricted to a vicinity of the position 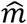, which by definition best agrees with activities from all modules, it was important to verify that the similarity of distances in light and dark trials was not an inevitable consequence of the methodology. Therefore, we introduced independent spatial shifts in the rate maps of all neurons that belong to the same module during the decoding process in a similar fashion as performed in the previous likelihood approach. We found that the mean distances *δ_i_* and Δ_*ij*_ increased dramatically as the magnitude of spatial shifts increased, indicating that these measured distances could have potentially been much higher than observed, and did not arise simply because of the selection of positions 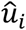 in proximity to 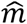 (Fig. 6c-d). Therefore, the preservation of these small distances in the dark, while the position 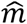 deviated substantially from the animal’s true position (Fig. 6e), is an explicit indication of tight coordination between the modules. As expected, identical spatial shifts did not affect *δ_i_* and Δ_*ij*_ even though they dramatically increased the MAE (faded traces in Fig. 6c-e). Similar results from additional datasets are shown in Fig. 6f-i and in Supplementary Fig. 9.

We finally tested whether module coordination remained tight in darkness, specifically during periods in which the multi-module readout 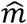 deviated substantially from the true position of the animal. To address this question, we focused on non-overlapping continuous segments of the dark recordings in which the MAE was particularly high, and on non-overlapping continuous segments within the same recording in which the MAE was particularly low (*Online Methods*). Even though the deviation of the internal representation from the true position was in the order of ~30 cm in the high-MAE segments (compared to order of ~5 cm in the low-MAE segments), the distances *δ_i_* remained small as in the low-MAE segments (Fig. 7a). Importantly, these distances *δ_i_* could have potentially been much higher, as demonstrated above (Fig. 6c-d). This result indicates that representations within individual modules did not accrue any additional, significant relative drifts even when the multi-module representation of position deviated substantially from the animal’s true position.

**Fig. 7:**
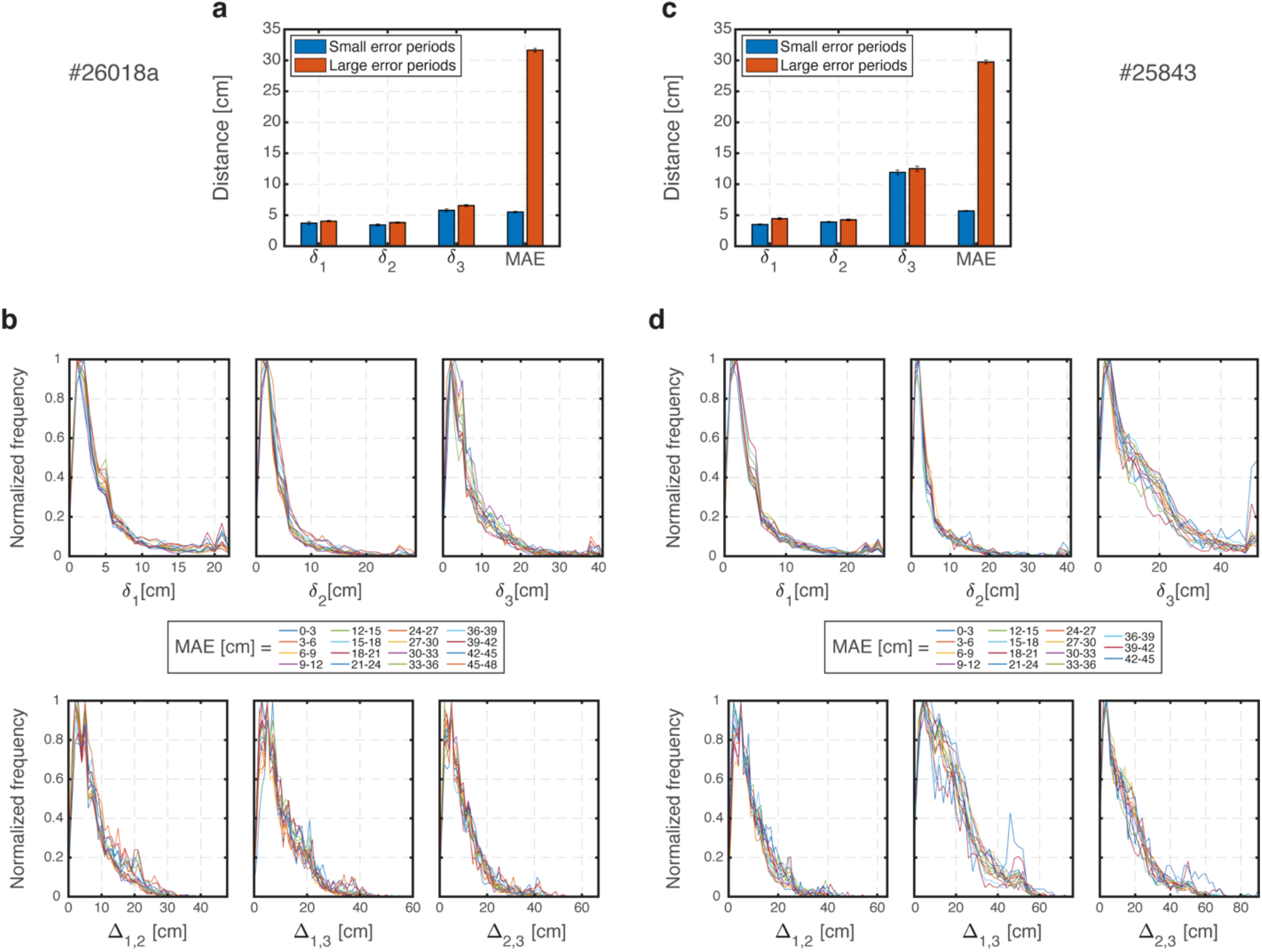
Module population activity patterns are tightly coordinated even during high MAE periods. **a,** The measured Mean Absolute Error (MAE) and distances *δ_i_* for dark and light during large- and small-MAE periods from a single recording session (#26018a). Error bars are ±SEM. **b,** Top: normalized distributions of the distances ő*i* for varying values of the MAE. Bottom: same as top but for distances Δ_*ij*_. **c-d,** Same as (a-b) but for another recording session (#25843). Note that outliers in (d) are traces with largest MAE.

Furthermore, we considered all time points from dark trials and looked on the joint distributions of *δ_i_* and the MAE, and on the joint distributions of Δ_*ij*_ and the MAE. We expected that if modules are coordinated then the conditioned distributions of *δ_i_* and Δ_*ij*_ will be nearly independent of the MAE, and in particular remain narrowly distributed even when the MAE is large. Fig. 7b demonstrates that, indeed, the distances *δ_i_* and Δ_*ij*_ were distributed nearly identically for different values of the MAE. This is yet another explicit indication that coordination between modules remains tight, even when the internal representation of position deviates from the true position of the animal. Similar results from additional datasets are shown in Fig.7c-d and in Supplementary Fig. 10.

## DISCUSSION

In contrast to the rigid relationships in activity of cells within a module, activity in different modules spans diverse phase combinations, allowing them to represent a large range of positions and environments. Nevertheless, here we showed that dynamically, the phases of different modules are coupled. Even when the internal representation of position in the MEC deviates substantially from the true position of the animal, updates to the phases remain coordinated across the different modules, thus maintaining a coherent representation of a two-dimensional trajectory.

The likelihood-based approach and the uni-module decoding approach both led to the conclusion, consistently across different animals and sessions, that the inter-module phases are dynamically coupled. Both methods also pointed to a small increase in the mismatch between modules in darkness compared to light conditions, indicating that sensory inputs help coordinate the states of different modules. The picture that emerges from these results is that when sensory inputs are poor or absent, small mismatches can develop in the phases of different modules, but internal brain mechanisms prevent these mismatches from accruing over time, thus maintaining a coordinated and coherent representation across the full grid cell population. It has been previously hypothesized that such internal mechanisms may exist (Burak, 2014; Welinder et al., 2008), possibly supported by recurrent synaptic connectivity within the MEC (Kang and Balasubramanian, 2019; Mosheiff and Burak, 2019), or by the reciprocal synaptic connectivity of the MEC with the hippocampus (Agmon and Burak, 2020; Sreenivasan and Fiete, 2011; Welinder et al., 2008). It will be of great interest to explore these underlying mechanisms in future studies, for example by testing whether inputs from the hippocampus are required in order to maintain phase coordination.

Despite a major reduction of sensory cues, their elimination was probably not flawless. Even though the arena was carefully cleaned during and between trials, leftover olfactory cues may have provided some spatial information. The rare encounters with the walls could also provide limited spatial information (Hardcastle et al., 2015), yet it is unlikely that absolute position could be inferred from such encounters as the arena was rotationally symmetric. Even if the elimination of sensory cues was imperfect, the disruption of the spatial rate maps and the increased MAE of decoded population activities in darkness (Fig. 2) showed that a large mismatch between internal representation and true position was typical under these conditions.

Previous studies that examined grid cell activity in mice in dark environments (Chen et al., 2016; Pérez-Escobar et al., 2016) have shown that even though individual cells’ periodic firing patterns lost their spatial stability, grid cells within a module preserved their pairwise spiking correlations, as observed in the light. However, these works did not test whether the coactivation patterns of distinct modules are coordinated during darkness. Due to the recording technique (classical tetrodes), simultaneously recorded cell pairs from distinct modules were rare, and the data collected did not enable analysis based on population decoding, as performed in this work.

Previous work (Stensola et al., 2012) pointed to a functional independence in the response of modules to an abrupt environmental deformation (Barry et al., 2007) in which the enclosure was compressed by moving one of the walls. A key finding was that this manipulation resulted in compression of the rate maps that occurred in some modules, but not in others. Thus, the distinct responses of different modules to the environmental deformation were indicative of functional independence in their dynamics. Nevertheless, the implications of this result for the dynamics of module coordination are not yet sufficiently clear: one possibility is that even shortly after the environmental deformation, grid cell firing remains anchored to position. In this case, the rates of phase updates, as a function of position, are modified compared to baseline conditions. Under this interpretation of the experiment, the dynamical coordination of modules is disrupted everywhere within the enclosure, possibly transiently. Alternatively, it has been suggested (Keinath et al., 2018; Ocko et al., 2018) that module phases are updated abruptly upon encounters with the walls, due to interactions with border cells. Under this interpretation of the experiment, rate maps are altered due to spatial shifts of the grid firing fields that depend on recent encounters with the walls. Yet, the phase update rates remain largely unmodified within the interior of the environment, and dynamical module coordination remains intact between encounters with the walls. Based on the tetrode recordings that were available in the deformation experiment (Stensola et al., 2012) it is difficult to conclusively distinguish between these possibilities. To do so, it will be beneficial to decode module phases from population activity patterns and analyze their joint dynamics, utilizing large numbers of simultaneously recorded grid cells from different modules.

The analysis in this work relied on the ability to simultaneously record spikes from dozens to hundreds of grid cells and our results demonstrate the power of this technique in elucidating dynamics within large neural circuits (Chaudhuri et al., 2019; Gallego et al., 2018; Gardner et al., 2021; Pfeiffer and Foster, 2013). With several dozens of cells from each module, decoding of phases from single modules was sufficient to obtain strong measures of coordination between the modules, based on statistics that were collected across long periods of motion. Future studies, with even larger numbers of simultaneously recorded cells, may enable more precise dynamical tracking of the states of individual modules over single trials. With such finer temporal and spatial resolution, it may be possible to characterize more precisely how the small mismatch that does exist between modules evolves over time, in relation to behavior or external stimuli. Such analysis may further elucidate the mechanisms that underlie coordination between attractor networks in the entorhinal cortex and the hippocampus.

## Online Methods

### Subjects

Experimental testing took place at the Kavli Institute for Systems Neuroscience, NTNU, Norway. Data were obtained from 4 male Long Evans rats (300-500 grams when implanted, at ages P 73-107 days old at day of recording). After weaning at three weeks, the rats were group-housed with their siblings until the implantation date. After implantation, each rat was housed alone in a large two-story enriched metal cage (95 x 63 x 61 cm). The rats were kept in temperature and humidity controlled rooms on a 12 hr light / 12 hr dark schedule. Experiments took place in the dark phase of the schedule. All procedures were performed in accordance with the Norwegian Animal Welfare Act and the European Convention for the Protection of Vertebrate Animals used for Experimental and Other Scientific Purposes.

### Electrode implantation surgery

The rats were implanted with single-shank 384-site Neuropixels probes (Jun et al., 2017) targeting the medial entorhinal cortex (MEC) in either one or both hemispheres. Rat #26018 was implanted only in the right hemisphere, while rats #25843 and #26820 were implanted bilaterally with prototype Neuropixels ‘phase 3A’ probes. Rat #26718 was implanted with a Neuropixels 1.0 probe in the right hemisphere. Before implantation, the rats were anaesthetized with isoflurane in an induction chamber and given subcutaneous injections of buprenorphine (Temgesic) and Meloxicam (Metacam). They were then fixed in a Kopf stereotaxic frame with continuous isoflurane administered through a mask. Local analgesic bupivacaine (Marcaine) was injected subcutaneously before making the incision. Craniotomies were drilled above the MEC area. The probes were inserted at a maximum depth of 5-6 mm from the brain surface, 4.4-4.6 mm lateral to the midline suture, 0.1-0.3 mm anterior to the transverse sinus, at angles between 25-26 degrees from the vertical plane, with the tip of the probe pointing in the anterior direction. A single jewellers screw was secured through the skull above the cerebellum and connected to the probe ground with an insulated silver wire. The implants were secured in place with dental adhesive (Optibond from Kerr) and Venus composite (Kulzer) and protected by fitting a modified falcon tube. Postoperative analgesia (meloxicam and buprenorphine) was administered during the surgical recovery period.

### Electrophysiological recordings

Electrophysiological signals were recorded with a Neuropixels acquisition system as described previously (Gardner et al., 2021; Jun et al., 2017). The spike band signal was recorded and amplified with a gain of 500, filtered to keep a bandwidth from 0.3 to 10 kHz and then digitized at 30 kHz on the probe circuit board. The signal was further multiplexed and transmitted to a Xilinx Kintex 7 FPGA board (‘phase 3A’) or a Neuropixels PXIe acquisition module (1.0) via a 5 m tether cable before being streamed via ethernet connection to a local computer. Rat #26018 had two recording sessions: recording session #26018a was performed 4 days before recording session #26018b, with partial overlap of recorded cells between the two sessions.

### Behavioural tracking

During recording, a rigid body with five retroreflective markers was attached to the rat’s implant and tracked with a 3D motion capture system (six OptiTrack Flex 13 cameras and Motive software) at ~120 Hz. To synchronise the timestamps of the two recording systems, randomized sequences of digital pulses generated by an Arduino microcontroller were sent to both the Neuropixels acquisition system as direct TTL input and to the OptiTrack system via infrared LEDs placed on the edge of the arena.

### Behavioural procedures

The rat’s movement was tracked as it moved freely in a circular open field arena. The recording arena was a 150 cm diameter, matt black plastic cylinder with 50 cm high walls and a matt black hard rubber floor, surrounded by floor-to-ceiling dark blue blackout curtains on all sides ~1m from the arena edge. Three additional layers of blackout curtains separated the recording arena from the part of the room with the recording computer. The same behavioural arena was used in both darkness and light recordings.

### Open-field foraging trials in darkness

Complete darkness was ensured by turning off all potential sources of light in the recording room. Light sources which could not be turned off were masked with aluminium foil and electrical tape and/or blackout curtains. Before starting the experiment, the arena and floors were thoroughly cleaned with soap water and dried. The rat’s Neuropixels probe was connected to the recording system outside the closed curtains before the final lights were shut off, and the rat was introduced to the recording arena at an arbitrary position and direction. For 50-60 minutes, the rat was left to freely explore the arena and forage small pieces of corn foam snack thrown into the arena during the trial by an experimenter wearing night-vision goggles (Armasight Nyx-7 pro). To avoid delivery of systematic orientational cues, the experimenter accessed and left the ring of curtains from random locations, and quickly dropped food pellets and removed excrement and urine using a paper towel while the rat was in a different location in the arena. Light conditions were not changed during these times.

### Open-field foraging task in light

After the open foraging task in darkness, while the rat was still foraging in the arena, or with a short break to untwist the Neuropixels tether cables, the experimenter turned on the light and continued the recording for another 30-60 minutes. During the light task, a single white textile cue card (~45 cm wide, ~150 cm high) hanging on the blue curtains outside the arena was visible from within the arena. The only light source in the light task was a single LED strip (6 m, 120 LEDs, 2800K color temp) placed as a uniform ~2 m diameter ring directly above the arena at a height of ~2.8 m, evenly illuminating the arena and ensuring no shadows were cast on the floor.

### Perfusion and histology

The rats were anaesthetized with isoflurane in an induction box and given a lethal injection of pentobarbital. When unresponsive, the rats were perfused transcardially with 0.9% saline, followed by 4% Formalin solution. The brain was extracted and stored in 4% Formalin solution for at least 24 hours before being sliced in 30 μm sagittal sections on a cryostat. The brain sections were stained with Cresyl Violet, and photomicrographs were taken through a Zeiss Axio Imager.

### Spike sorting and single-unit selection

Spike sorting was performed with a version of KiloSort 2.5 (Steinmetz et al., 2021), optimised for MEC/PaS recordings as described in (Gardner et al., 2021), including manual supervision of cluster split and merge processes. Single units were excluded from further analysis if more than 1% of intervals in their interspike interval distribution were shorter than 2 ms or if they had less than 500 total spikes in the light task.

### Module classification

Grid cell and module classification was done by vectorizing the spatial autocorrelation from the rate map of every cell and adding them as feature columns in a matrix used as input to the UMAP (Uniform Manifold Approximation and Projection) dimensionality reduction algorithm (McInnes et al., 2018), before DBSCAN (Ester et al., 1996) was used to assign cluster identities to the resulting 2D point clouds, as in (Gardner et al., 2021) (Fig. 1e-f and Supplementary Fig. 2a-c). Briefly, for each recorded cell, rate maps were generated by dividing the arena into 8-10 cm bins and counting the number of spikes within each bin divided by the time spent in that bin. Autocorrelograms of the rate maps were calculated, and values from the bins in a circular area within a 3-bin radius from the center bin were removed along with the bins outside a radius defined by the edge of the matrix. The autocorrelograms were then vectorized and used as features in UMAP used to project the values down to a point cloud in 2 dimensions. DBSCAN was used to cluster the points, which yielded a single large cluster with non-grid cells and single clusters for each module (identified by a clear grid pattern and high gridness score in the mean autocorrelogram of each cluster); grid modules could be ordered by the grid spacing and orientation calculated from the mean autocorrelogram of each cluster. Of all recorded cells, only those from clusters with a clear grid pattern in the mean autocorrelogram, and a validity index (see clustering validation section) close to 1.0 were used for further analysis.

### Clustering validation

The grid cell and module classification results were validated by calculating a density-based clustering validation (DBCV) index (Moulavi et al., 2014) for each DBSCAN-assigned cluster identity in the 2D UMAP point cloud (Fig. 1e-f and Supplementary Fig. 2a-c). The DBCV index has a range of −1 to 1, where a cluster gets a positive value if the lowest density region inside the cluster is higher than the highest density in the region that separates it from other clusters (Supplementary Fig. 11).

### Rate map analysis

The firing rate *λ_i_* [Hz] of grid cell *i* at each position 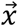 in the arena was generated as follows:

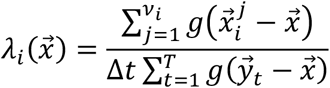

where *v_i_* is the total number of spikes emitted by neuron *i* and 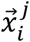 is the animal’s position when spike *j* was emitted. The position of the animal at time *t* is denoted by 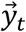, and *g* is a two-dimensional Gaussian kernel with diagonal covariance matrix Σ_*ii*_ = 25 cm^2^. Spike trains and tracking data were binned at 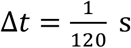 resolution, and only time bins where the animal was moving at a speed greater or equal to 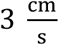 were used for spatial analyses.

### Gridness score

The gridness score was computed to measure the degree of hexagonal spatial periodicity, as in (Langston et al., 2010). For each cell, an autocorrelogram was calculated from its rate map and rotated in five steps of 30 degrees, correlating each rotated matrix with the original autocorrelation in the following manner: First, the values correlated were restricted to a ring of bin indexes around the center peak of the autocorrelogram; then, this ring of bins was expanded stepwise until its outer edge reached the edge of the autocorrelogram matrix. For each step, a score was calculated as the difference between the lowest correlation at [60, 120] degrees and the highest correlation at [30, 90, 150] degrees. The gridness score was taken as the mean of the three scores surrounding and including the step with the maximum score, resulting in a theoretical score range of [-2, 2].

### Information content

The spatial information content [bits/spike] (Skaggs et al., 1996) of neuron *i* is defined as

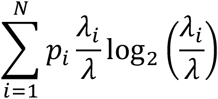

where *λ_i_* is the unit’s mean firing rate in the *i*-th bin of the rate map, *λ* is the overall mean firing rate and *p_i_* is the probability of the animal being in the *i*-th bin (time spent in the *i*-th bin divided by the duration of recording).

### Pairwise correlations

Spike trains were binned at 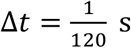 resolution and smoothed using a Gaussian kernel with *σ* = 50 ms. Pearson correlation coefficients were then calculated for pairs of spike trains from simultaneously recorded grid cells.

In Figure 3b,d neurons were divided into ten equally sized groups with equal distributions of neurons from each module and cross-correlations were calculated across inter- and intra-module pairs that belong to the same group to obtain independent evaluations. Cross-correlations were calculated by iteratively lagging one of the spike trains relative to the other. Note that since both negative and positive correlations are meaningful as positive ones, we averaged the absolute magnitude of the correlation over pairs within each of the independent groups.

In Figure 3c,e and Supplementary Figure 5b, the Pearson correlation coefficients were calculated for all possible inter- and intra-module pairs without lagging any of the spike trains.

### Markov decoder

The Markov decoder updates its posterior likelihood for position *r* as follows:

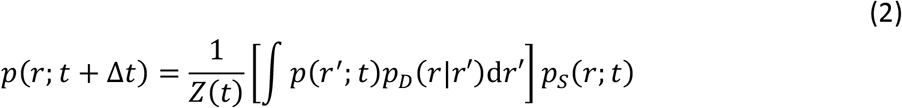

where *p_D_*(*r|r′*) describes the animal’s probability to run from location *r′* to location *r* during time interval Δ*t*. The term *p_s_*(*r; t*) is the probability for all the neurons to emit the observed spikes within the time interval Δ*t*., given the position *r*. The posterior likelihood is iteratively normalized by *Z*(*t*).

Explicitly, *p_D_*(*r|r′*) is a two-dimensional Gaussian distribution centered around position *r′* with diagonal covariance matrix Σ_*ii*_ = 4 cm^2^, and 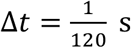.

The posterior extracted from spiking activity was evaluated assuming independent Poisson firing, namely

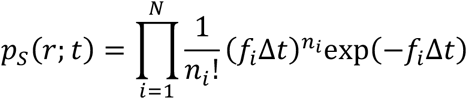

where *f_i_*(*r*) is the tuning curve and *n_i_* is the spike count of the *i*’th neuron during the time interval Δ*t*.

Finally, the estimate of position is the maximum likelihood estimate

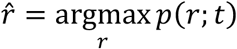

### Likelihood of simultaneously recorded spike trains

The probability for observed spike trains *p*(***S**_t_*) is given in Eq. 1. By exploiting the Markov decoder’s properties, the likelihood for simultaneous observed spikes turns out to be simply proportional to the multiplication of its iterative normalization factors presented above,

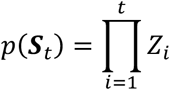

A detailed analytical derivation is given in the *Supplementory informotion*.

We present in Figure 4 and 5 the average log likelihood per time unit defined as

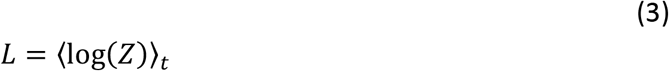

Since decoding was Markovian we decoded all the data, but only time bins where the animal was moving at a speed greater or equal to 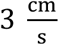 were used for further analyses.

### Rate-adjusted likelihood

In order to evaluate the likelihood using the Markov decoder and faithfully compare this quantity across light and dark conditions, it is necessary to make concrete assumptions on the neural tuning curves. Since individual grid cells differed in their firing rates in these two conditions, it was necessary to analyze how the firing rate influences the likelihood, and then compensate for this influence. Counter intuitively, the evaluated likelihood decreases as the number of total spikes in the recording increases (see *Supplementory informotion* for an analytical derivation which elucidates this finding).

Our assumption is that neurons maintain the same underlying structure of tuning curves in light and dark conditions, up to a scaling factor that adjusts the firing rate (see also *Supplementory Informotion*). Therefore, we matched the mean firing rates by randomly omitting spikes in the condition in which the firing rate was higher, yielding spike trains from each neuron with the same mean firing rate in the dark and light conditions. This was done separately for time bins above speed greater or equal to 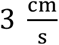 since these time bins were used for further analysis.

### Spatial Shifts

For each module, a fixed spatial shift in each of the two spatial dimensions was chosen independently and at random from the range [−*α, α*]. The variable *α* is plotted in the ‘spatial shift’ axis in figures throughout this article. Rate maps of all neurons that belong to the same module were shifted according to corresponding random shifts and were then trimmed at the arena boundaries in their original positions, thus producing shifted rate maps. Firing rates at new positions that were included within the arena boundaries only after the shift but were outside of the arena boundaries before the shift were set to zero. Identical spatial shifts were applied in a similar procedure, but only a single set of two-dimensional shifts were chosen at random and applied to the rate maps of all neurons as described above, regardless of the module they belong to.

### Kernel decoder

The kernel decoder updates its posterior likelihood for position *r* based on recent emitted spikes, weighted exponentially (Mosheiff et al., 2017). It is straightforward to express the log likelihood of spike counts *v_i_*, observed within a temporal window of duration Δ*t*, as a function of the position *r*:

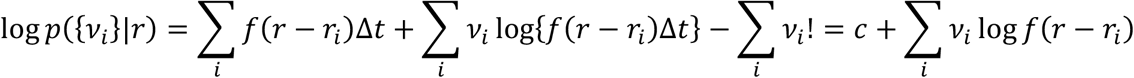

where *c* is a constant that does not depend on *r*. The term Σ_*i*_ *f*(*r* – *r_i_*) contributes only to this constant because of the assumption of dense, transnationally invariant receptive fields with uniform distribution. A maximum likelihood estimator for *r* (assuming uniform prior) will chose

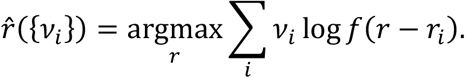

The spikes from recent history are weighted with a temporal kernel *h*(*t*). Thus, we generalise *v_i_* to:

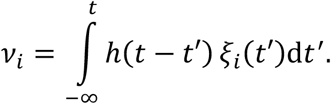

Here, *ξ_i_*(*t*) is a series of delta functions that represents the spike from neuron *i*. The counting of spikes within a temporal window of duration Δ*t* under this definition is achieved by setting *h*(*t*) to

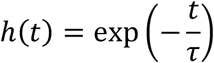

where we used *τ* = 100 ms.

Only time bins where the animal was moving at a speed greater or equal to 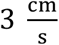 were used for analyses.

### Uni-module decoding

As expected, when decoding activity from single modules the posterior was approximately periodic. To remove ambiguity, the decoded position 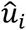 of each module *i* was defined as the position that maximized the posterior within a circular area. The circular area had a diameter equal to ~90% of the corresponding module spacing and was centered around the multi-module position 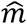 (Fig. 6a). Thus, only a single blob was included within the circular area around the multi-module represented position 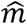.

### High and low Mean Absolute Error (MAE) segments

In Fig. 7a,c the MAE was temporally smoothed using a Gaussian kernel with *σ* = 50 ms (sMAE). Segments with particularly low MAE are defined as continuous non-overlapping intervals spanning at least 1 s with a maximal sMAE of 10 cm. Segments with particularly high MAE are defined as continuous non-overlapping intervals spanning at least 1 s with a minimal sMAE of 20 cm, and a maximal sMAE which was determined as follows: the maximal sMAE was chosen as the value corresponding to 80% of the cumulative distribution of the MAE. Time points with higher MAE were discarded from the analysis in accordance with the cutoff used in Fig. 7b,d (largest MAE value) due to sparseness of the joint distribution with the distance between 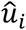 and 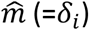. Recording session #25843 had 257 high-error segments and 305 low-error segments. Recording session #26018a had 426 high-error segments and 126 low-error segments. Recording session #26018b had 460 high-error segments and 156 low-error segments. Recording session #26820 had 384 high-error segments and 92 low-error segments.

### SEM of correlated time series

Whenever the standard error of the mean (SEM) of a single time series signal (*S*) was evaluated, correlations were taken into account by updating the signal’s variance based on the auto-correaltion function (*ACF*(*S*)).

For an independent signal the variance is simply *ACF*_0_. However, for a temporally correlated stationary signal, the actual variance is written as

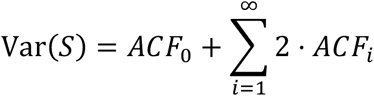

where in practice the cutoff point of the sum remains to be determined. We trimmed the sum at a point corresponding to an autocorrelation value satisfying *ACF_i_* ≤ 0.15 · *ACF*_0_. Finally, the SEM is defined as 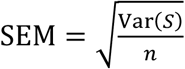 where *n* is the total number of points in the signal.

### Data availability

Data is available upon request.

### Code availability

Code is available upon request.

## Acknowledgements

Y.B. is the incumbent of the William N. Skirball Chair in Neurophysics. This study was supported by a Synergy Grant to Y.B. and E.I.M. from the European Research Council (‘KILONEURONS’, Grant Agreement No. 951319); by grants to Y.B. from the Israel Science Foundation (Grant Nos. 1319/13, 1978/13, and 1745/18); by a grant to Y.B. from the German-Israeli Foundation for Scientific Research and Development; a FRIPRO grant to E.I.M., a Centre of Excellence grant to M.-B.M. and E.I.M., and a National Infrastructure grant to E.I.M. and M.-B.M, all from the Research Council of Norway (FRIPRO grant number 286225; Centre of Neural Computation, grant number 223262; NORBRAIN, grant number 295721); the Kavli Foundation (M.-B.M. and E.I.M.); and a direct contribution to M.-B.M. and E.I.M. from the Ministry of Education and Research of Norway. Y.B. acknowledges support from the Gatsby Charitable Foundation. The authors are grateful to Christine Lykken for performing the Neuropixels implantation on one of the animals.

## Author contributions

E.I.M. and Y.B. conceived the study; T.W., M.-B.M. and E.I.M. designed the experiments; T.W., R.J.G., V.A.N. and A.N. performed the experiments (surgeries, recordings, spike sorting); R.J.G. developed Neuropixels data analysis pipelines; H.A. and Y.B. conceived and developed the theory and the computational methodology; T.W. and H.A. performed single cell analyses; T.W., R.J.G. and V.A.N. performed module clustering (UMAP); H.A. performed the simulations; H.A. analyzed module coordination; H.A., E.I.M. and Y.B. interpreted module coordination results with help from all authors; T.W. and H.A. visualized the data; H.A., E.I.M. and Y.B. wrote the paper with contributions from T.W. and inputs from all authors; R.J.G., M.-B.M., E.I.M. and Y.B. supervised the project; M.-B.M., E.I.M. and Y.B. obtained funding.

---

**Supplementary Fig. 1:**
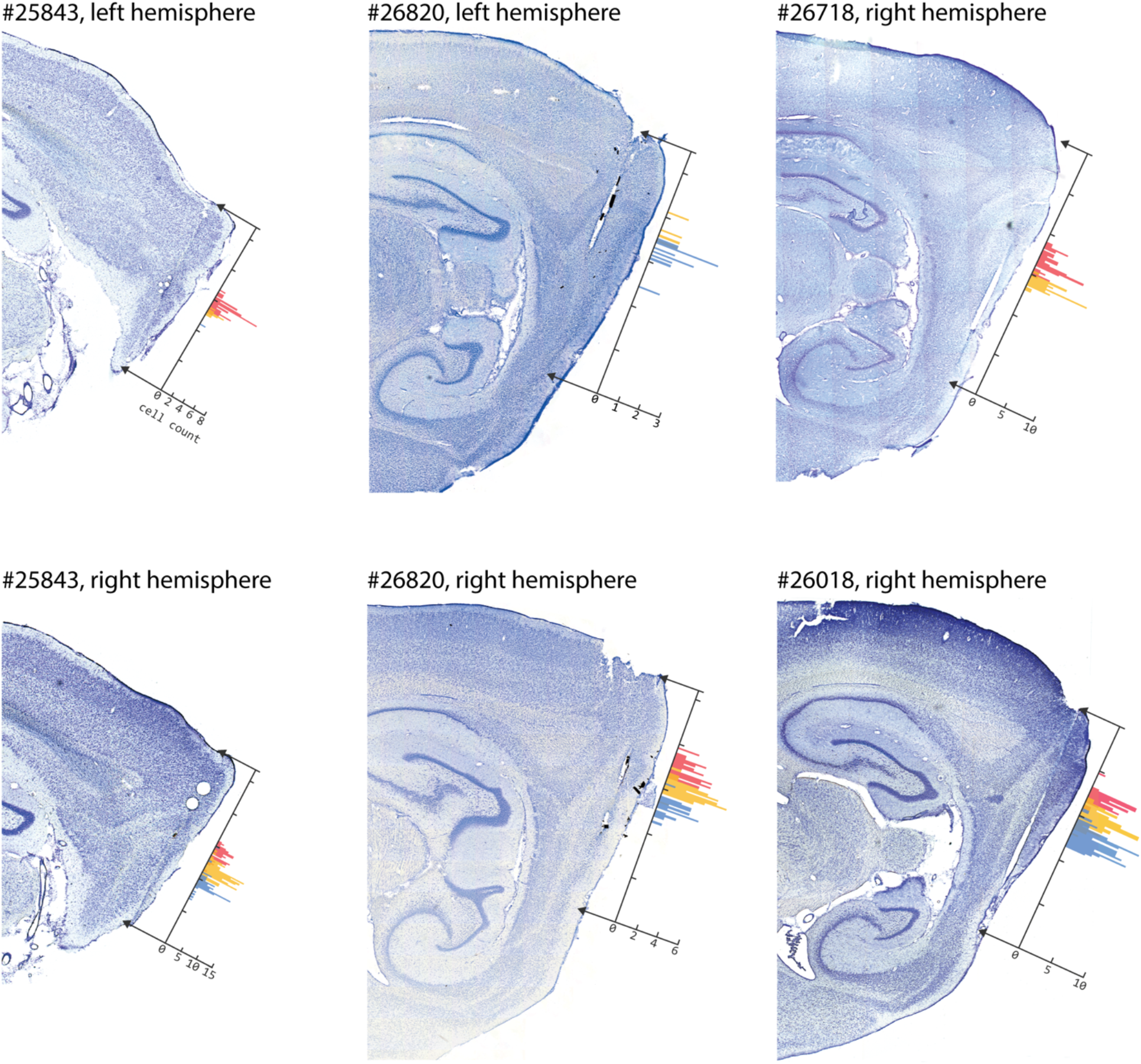
Histology and estimated recording sites. One cresyl violet-stained sagittal section is shown for each rat’s neuropixels probe, showing the probe track left in the brain tissue. Estimated entering sites in the brain as well as probe tip locations are marked with arrows. The histogram shows the grid cell count across dorso-ventral recording depths from different modules (color coded). The distance between two adjacent ticks along the probe shank axis corresponds to 1 mm.

**Supplementary Fig. 2:**
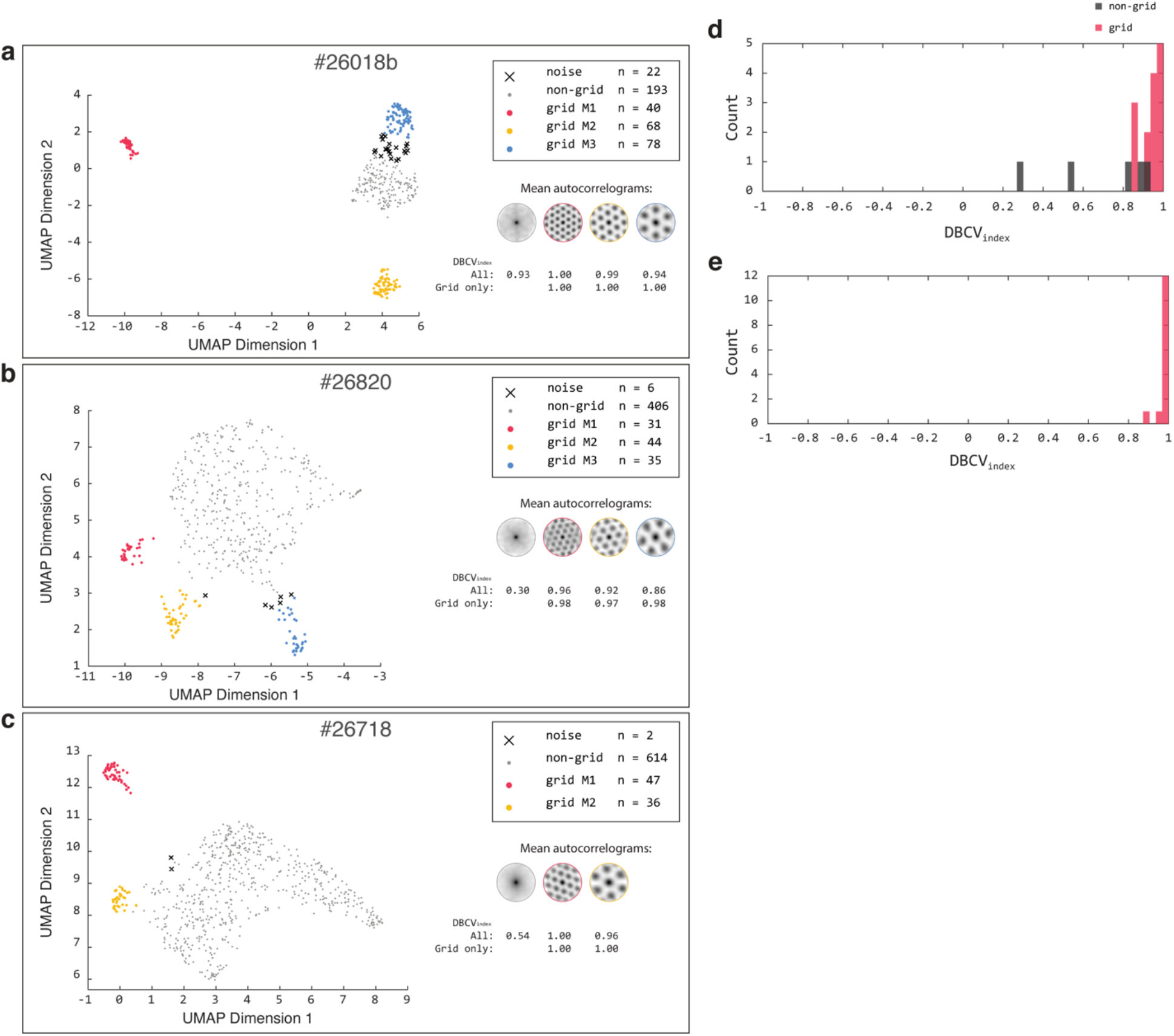
Grid cell classification by UMAP-DBSCAN. **a-c,** Same as Fig. 1e-f but for three additional recording sessions. **d,** Distribution of validity index with non-grid cluster included for all rats and recordings. **e,** Distribution of validity index with non-grid cluster excluded for all rats and recordings.

**Supplementary Fig. 3:**
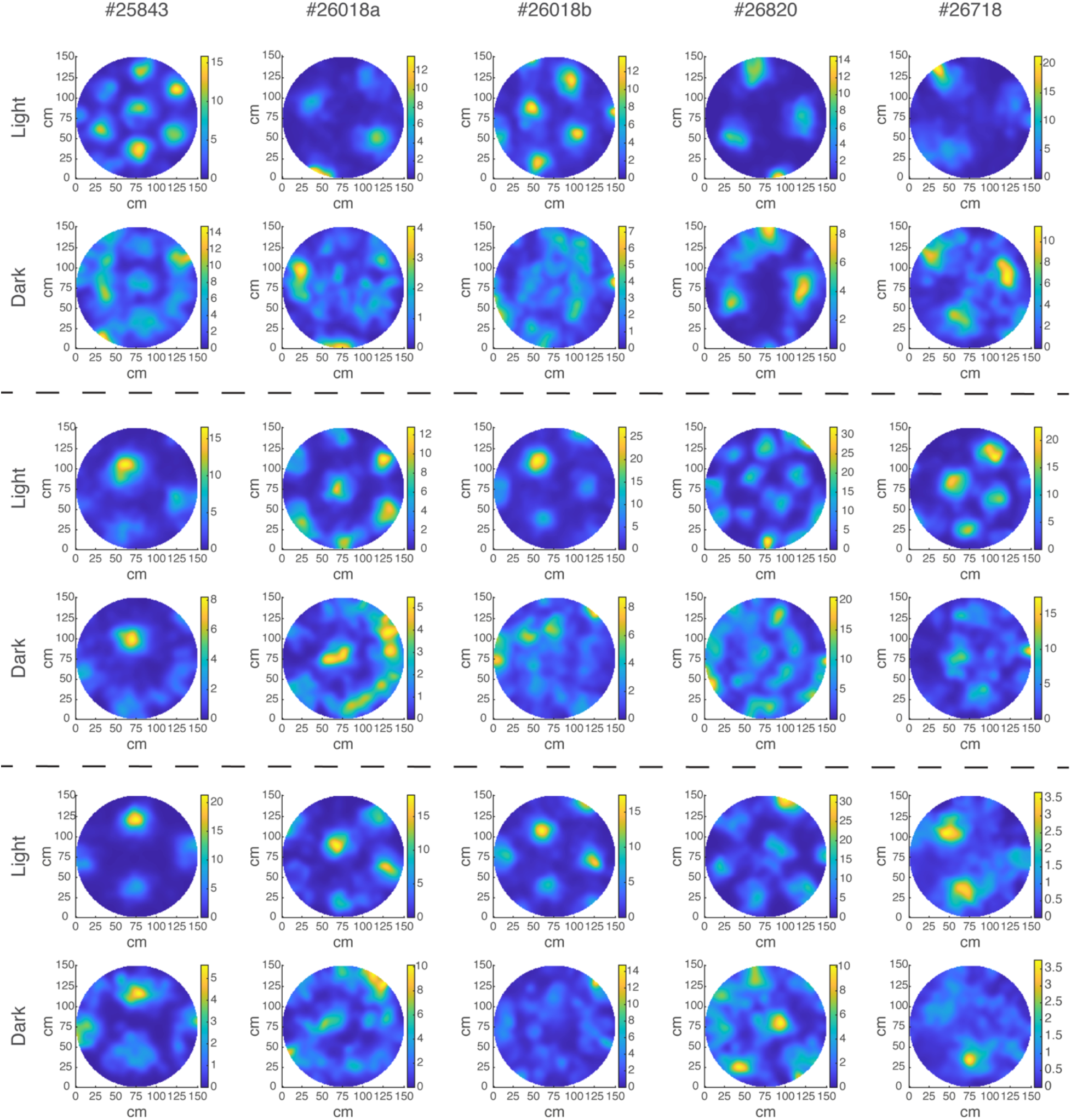
Examples of individual grid cell rate maps from light and dark. Same as Fig. 2a, but for another three examples from each recording session.

**Supplementary Fig. 4:**
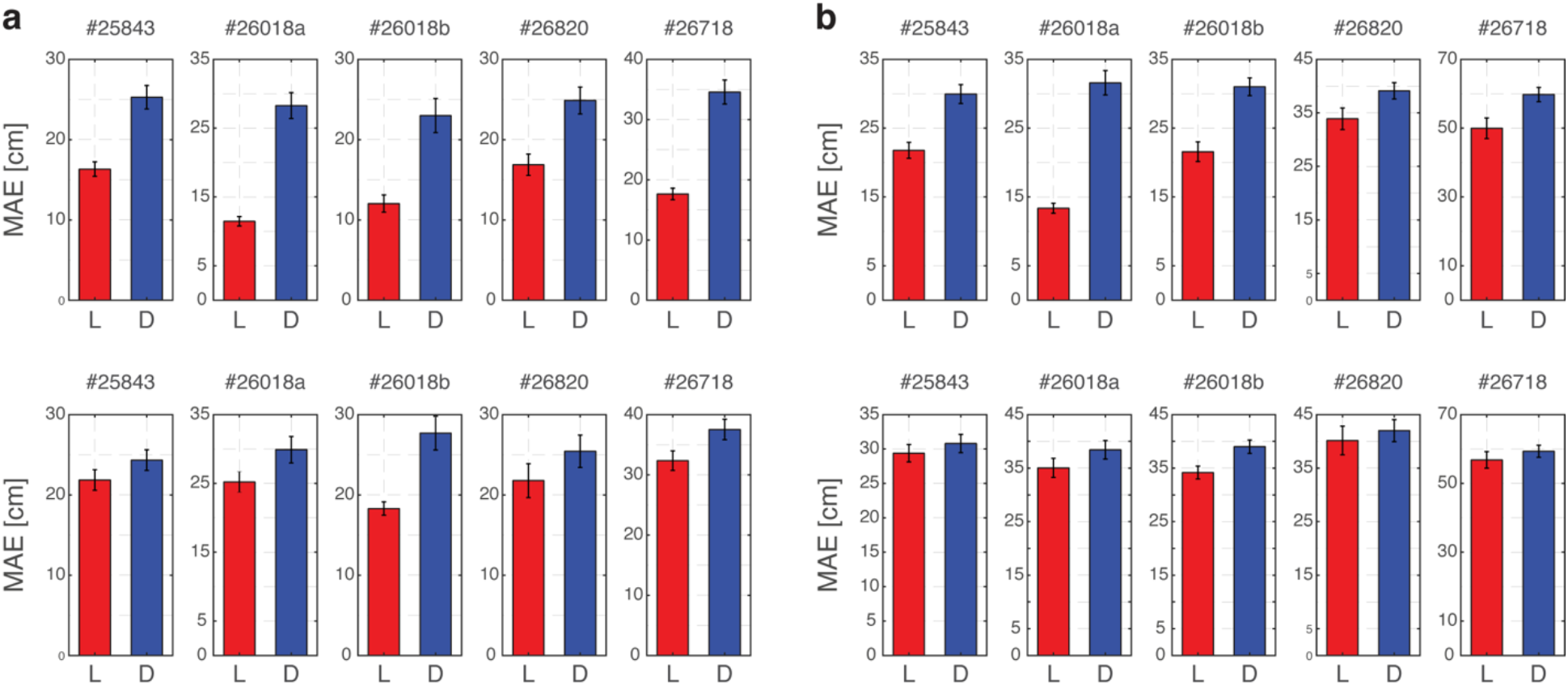
Decoding performance does not improve even when using dark generated rate maps. **a,** Top: Mean Absolute Error (MAE) of the Markov decoder applied on the light and dark recorded spiking activity for all cells from single recording sessions and when rate maps were constructed only from one half of the light data (and used to decode the other half of data). Bottom: same as top, but when rate maps were constructed only from one half of the dark data (and used to decode the other half of data). As expected, using the dark-generated rate maps leads to an increase of the MAE both in light and dark conditions, relative to decoding using light-generated rate maps. Importantly, the MAE of decoded position in the dark recording sessions is still larger than in light sessions, even when dark-generated maps are used. Error bars are ±SEM. **b,** Same as (a) but for the kernel decoder.

**Supplementary Fig. 5:**
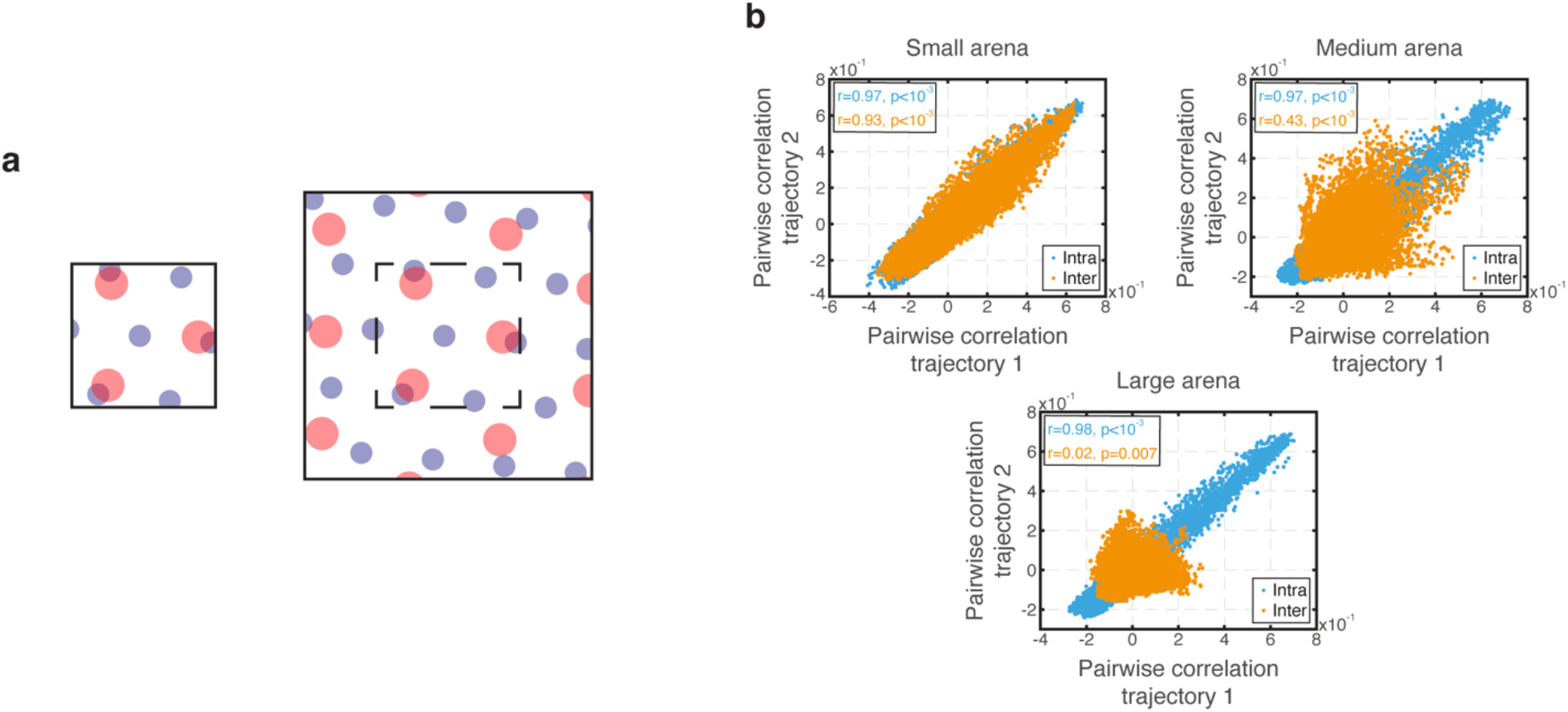
Inter-module pairwise correlations diminish with increased environment size. **a,** Schematic illustration showing spatial tuning curves of two inter-module grid cells (blue and red). The mean correlation between the tuning curves is higher when the environment is small (left) than when it is large (right). In larger environments, the spiking correlations are expected to be more narrowly distributed around zero. **b,** Pairwise correlations of all possible intra- (cyan) and inter- (orange) module pairs from two simulated trajectories in small (radius=30 cm), medium (radius=60 cm) and large (radius=150 cm) circular arenas. Correlation coefficients and p-values are specified in the insets. As expected, inter-module spiking correlations became narrowly distributed around zero with an increase in the size of the environment, whereas intramodule spiking correlations remained unaffected. Simulated grid cells share the spacings and module allocation as recorded in session #26018b and emit Poisson spikes which are determined by their idealized tuning curves.

**Supplementary Fig. 6:**
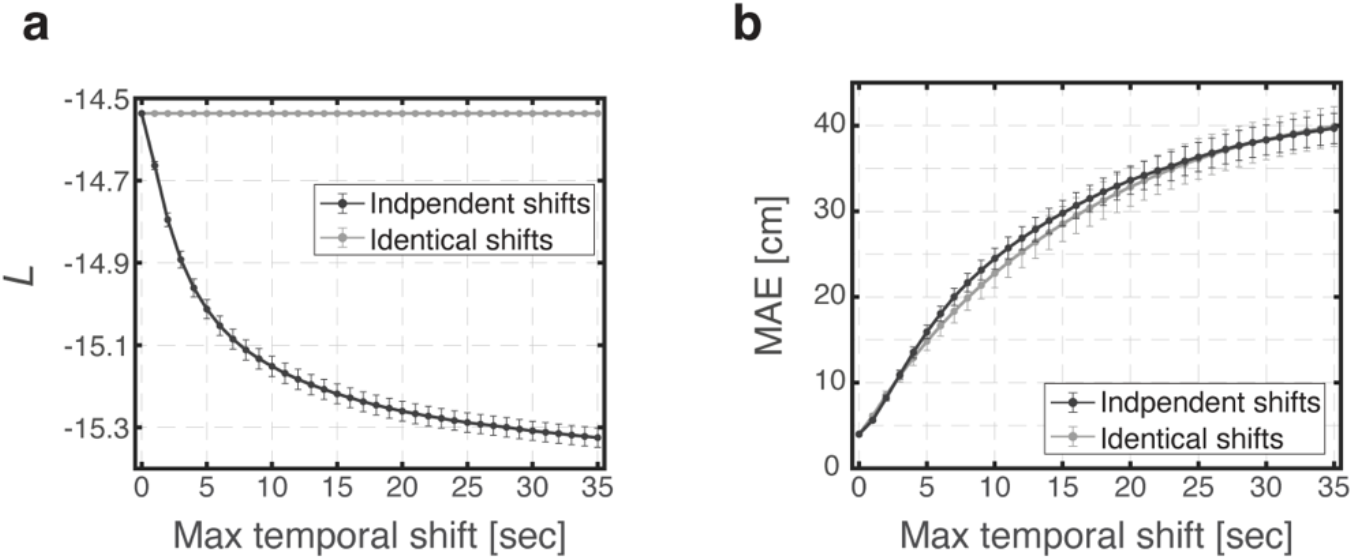
Likelihood slightly decreases under identical spatial shifts due to boundary conditions. In order to demonstrate that the slight decrease in the likelihood when using identical spatial shifts is due to boundary conditions, temporal shifts were applied to the same data as in Fig. 4 b-c. Independent temporal shifts were applied by shifting the timing of spike trains of all neurons that belong to the same module, and identical temporal shifts were applied in a similar fashion but identically for all neurons regardless of the module they belong to. Rate maps were unaffected during this procedure thus their boundaries were not trimmed as in the spatial shift procedure. **a,** Likelihood of simulated Poisson spikes using measured rate maps and recorded light trajectory from session #26018b for varying magnitudes of temporal shifts. When shifts are applied independently for each module, the likelihood decreases significantly while remaining precisely fixed when temporal shifts are identical. Error bars are ±SEM. **b,** The Mean Absolute Error (MAE) increases significantly both for independent and for identical temporal shifts as their magnitude increases. Error bars are ±SEM.

**Supplementary Fig. 7:**
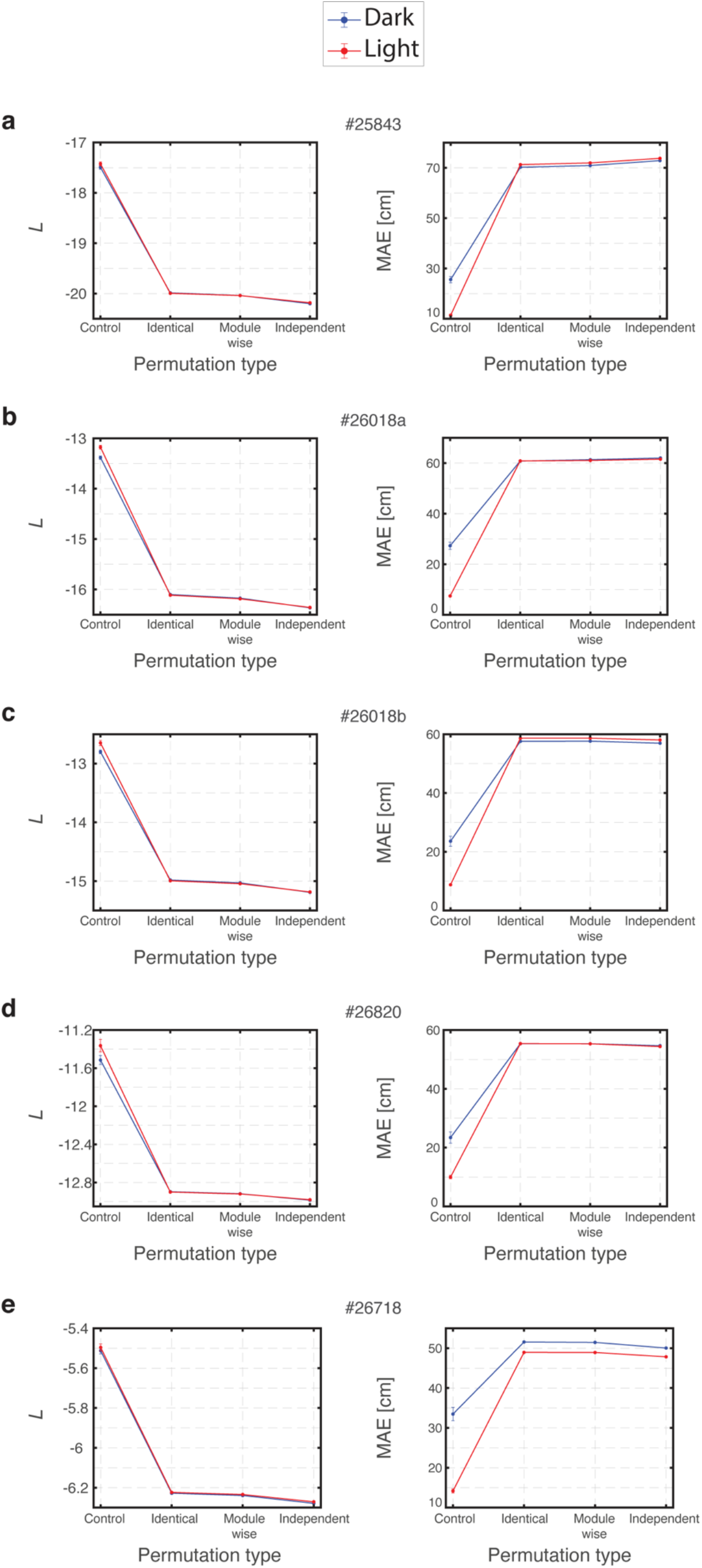
The temporal structure of simultaneously recorded spike trains, and not simply their mean firing rate, is necessary to account for the likelihood and MAE results. Three types of permutations, which preserve the mean firing rates, were applied to the simultaneous recorded spike trains. ‘Identical’ describes a permutation type in which the exact permutation was applied to all neurons, ‘Module-wise’ describes a permutation type in which identical permutations were applied but only to neurons that belong to the same module, and ‘Independent’ describes a permutation type in which an independent permutation was applied to each neuron. **a-e,** Likelihood of simultaneous recorded spike trains (left) and corresponding Mean Absolute Error (MAE, right) from dark and light trials for the different permutation types and for all recording sessions. Error bars are ±SEM.

**Supplementary Fig. 8:**
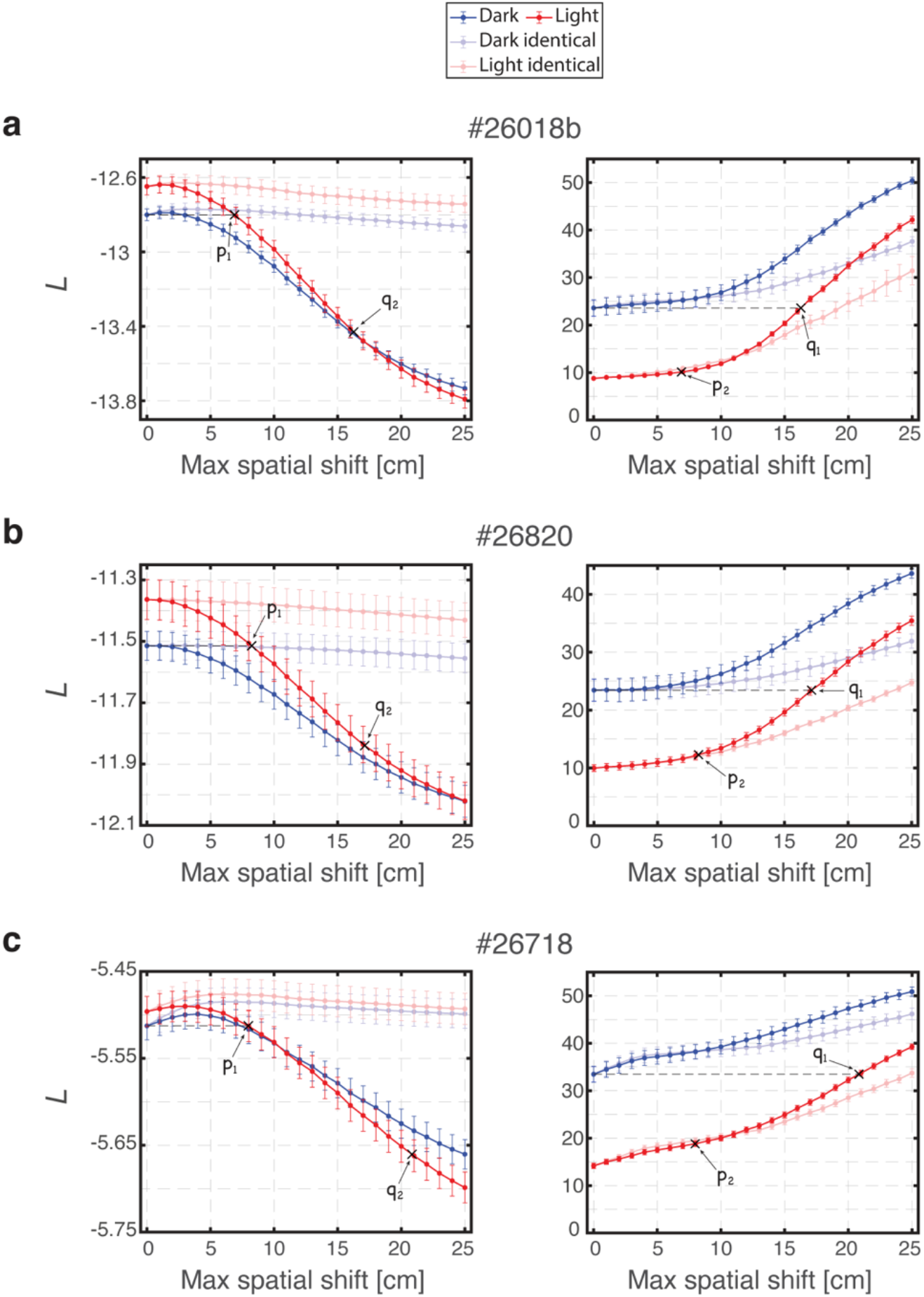
**a-c,** Same as Fig. 5c, for three additional recording sessions.

**Supplementary Fig. 9:**
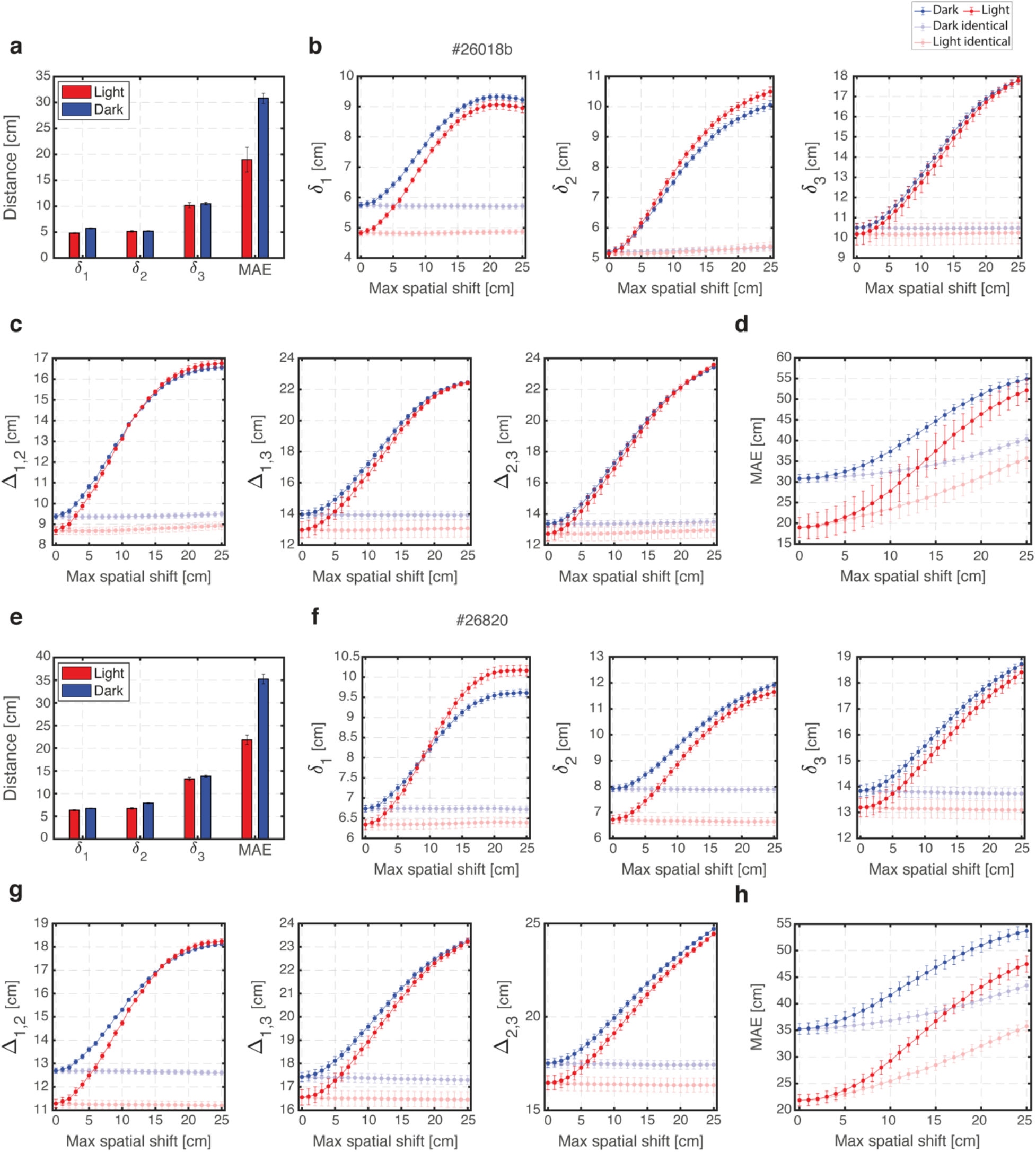
**a-d** and **e-h,** Same as Fig. 6b-e for two additional recording sessions (#26018b and #26820) in which recordings were obtained from three modules. Note that even though recordings from three modules were available in these datasets, the numbers of simultaneously recorded grid cells were relatively small (especially in recording session #26820; Table 1), leading to inaccurate decoding.

**Supplementary Fig. 10:**
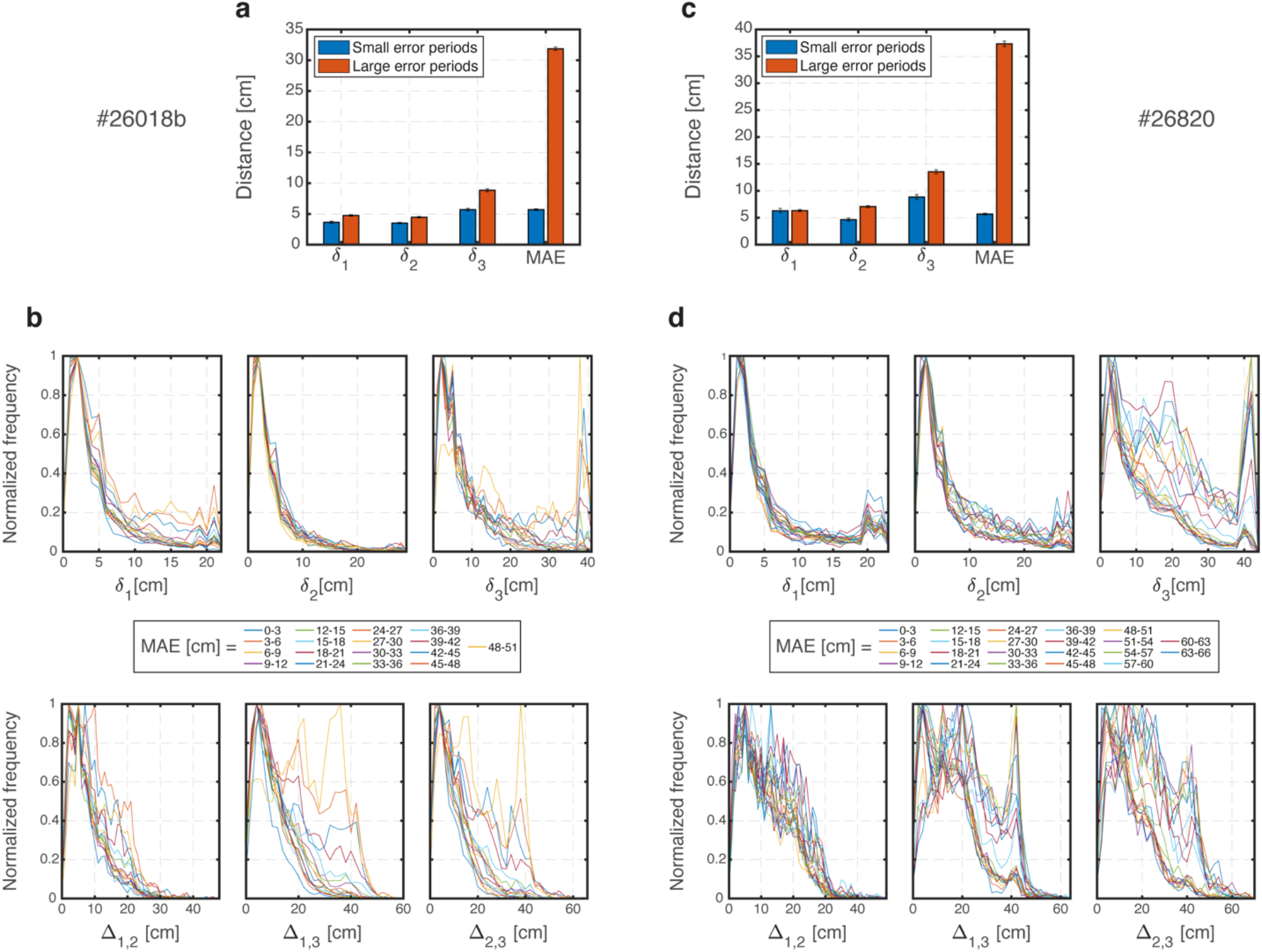
**a-b** and **c-d,** Same as Fig. 7a-b, for two additional recording sessions (#26018b and #26820) in which recordings were obtained from three modules. Note that even though recordings from three modules were available in these datasets, the numbers of simultaneously recorded grid cells were relatively small (especially in recording session #26820; Table 1), leading to inaccurate decoding.

**Supplementary Fig. 11:**
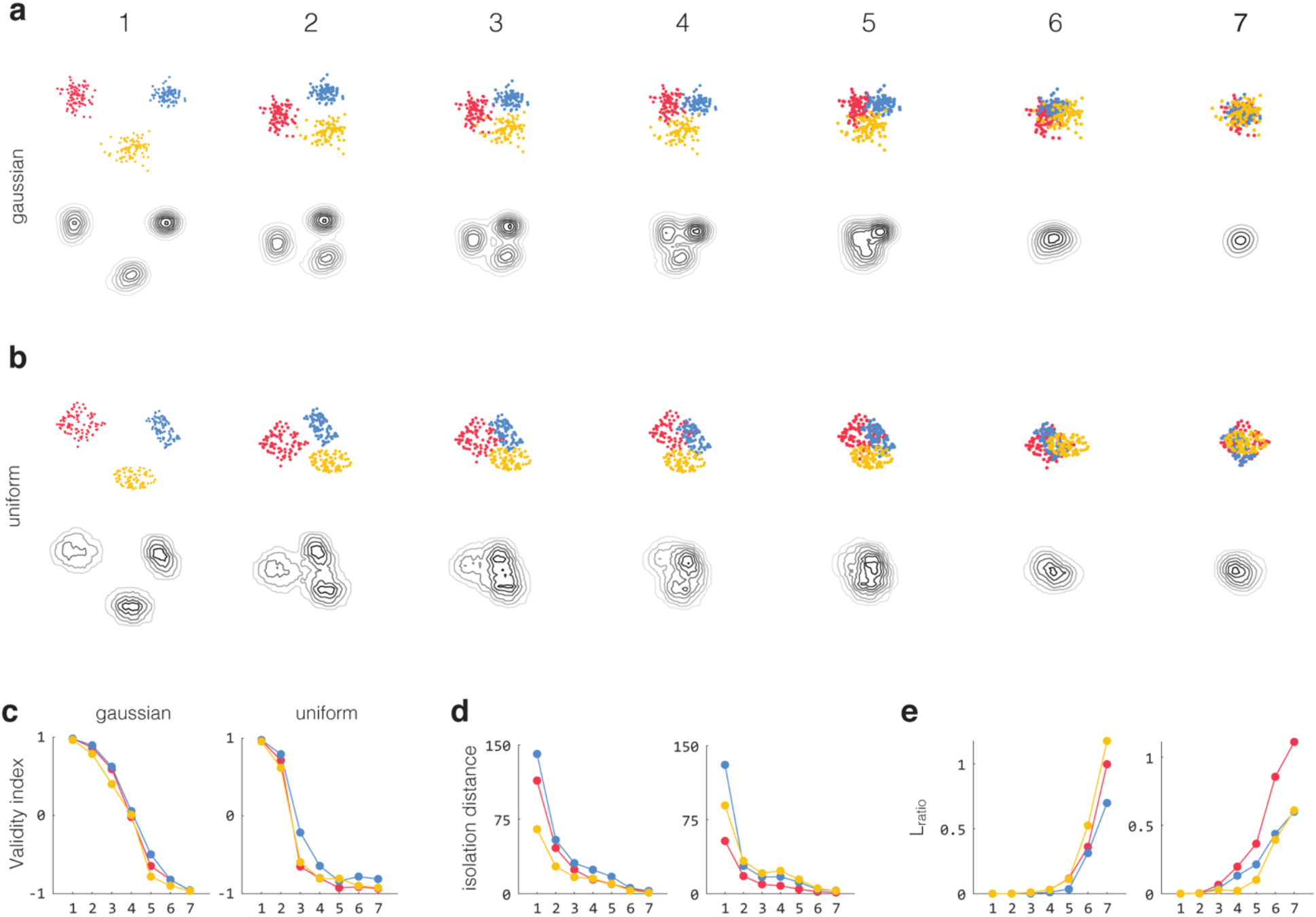
Clustering quality validation: examples of DBCV index on varying clustering quality. Synthetic data points in clusters with varying degree of separation, either random two-dimensional gaussian distributions (a) or random uniform distributions with different shapes (b). All plots have the same scale. **a-b,** Top: synthetic data scatterplot with colour coded cluster assignment. Bottom: contour plots illustrate the density. **c,** Density based clustering validity (DBCV) index of the clusters in (a, left) and (b, right), with colour corresponding to cluster. Well separated clusters have a DBCV index above zero. **d-e,** We include the isolation distance and L-Ratio measures (Scmitzer-Tobert 2005) commonly used for comparison of clusters of tetrode-recorded spikes from the hippocampus. The two measures were developed for measuring clustering quality for spike sorting tetrode data and assume the clusters form a gaussian distribution in feature space; they might not be ideal for quantifying density-based clustering results in UMAP space.

**Supplementary Fig. 12:**
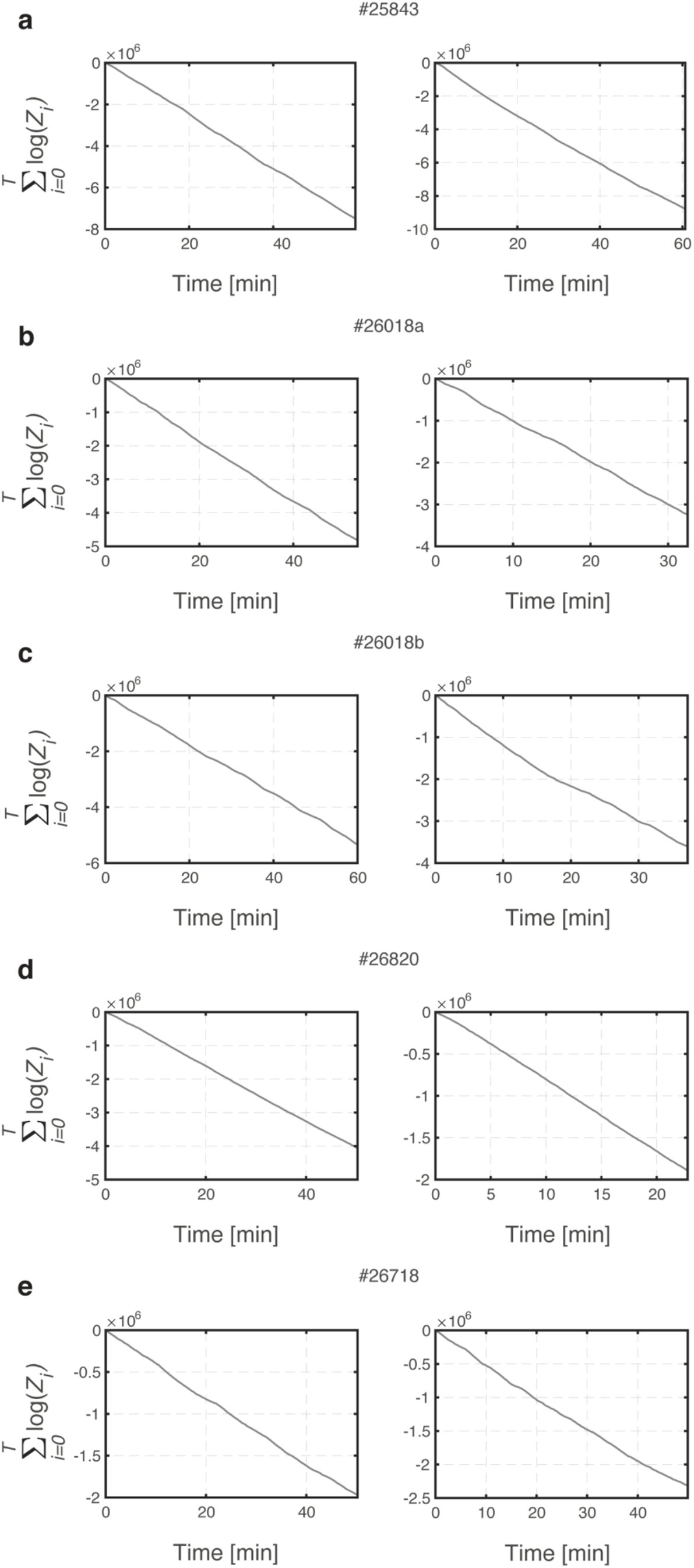
Log likelihood versus time. **a-e,** Cumulative sum over time of log[*p*(***S**_t_*)] in dark (left) and in light (right) trials from all recording sessions. Linear dependence (R^2^>0.99 for both light and dark and across all sessions, fit not shown) is evident, indicating a steady accumulation of the log likelihood per time bin.

**Supplementary Fig. 13:**
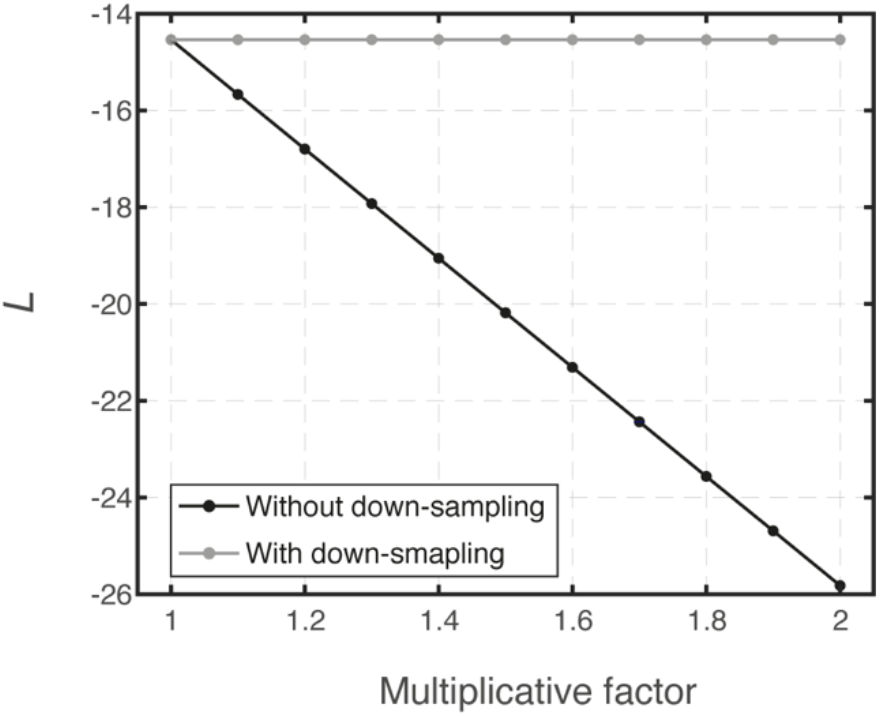
Likelihood dependence on the mean firing rate. The likelihood *L* was evaluated when using rate maps and light trajectory from recording session #26018b but with simulated Poisson spikes. Rate maps were first scaled by a multiplicative factor (x axis) thus leading to a constant decrease in the likelihood as the firing rate increases (black trace, as expected by Eq. S12). However, when the spike trains that were generated using the scaled rate maps were randomly down-sampled to match their original mean firing rate, the original likelihood value was precisely restored (gray trace).

## Supplementary information

### a. Likelihood of simultaneously recorded spike trains

In this subsection we mathematically derive the likelihood of simultaneously recorded spike trains, independently from the animal’s true position. To address the inter-module coordination question, we sought to derive a measurement which can quantify the coherence of the simultaneously recorded spike trains even if the represented position in the brain is dissociated from the animal’s true position.

Denote by 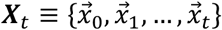 a particular realizable two-dimensional trajectory up to time *t*, and by *p*(***X**_t_*) the probability for the trajectory ***X**_t_* to be realized. Since the prior on the trajectory is Markovian, *p*(***X**_t_*) satisfies

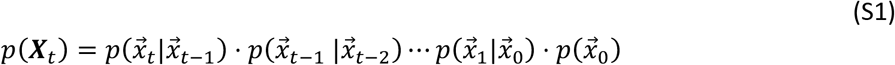

where 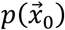 is the probability of the initial position.

The probability to observe the simultaneous recorded spike trains, averaged over all possible trajectories up to time *t*, is written as

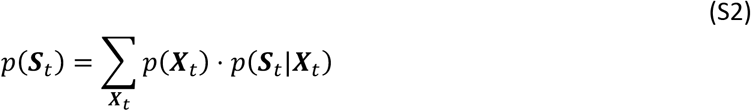

where the sum is over all possible trajectories, weighted by their corresponding priors *p*(***X**_t_*). Extending Eq. S2 for the consecutive time step *t* + 1,

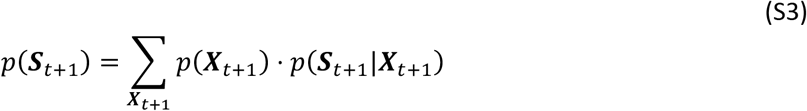

Due to the Markov properties 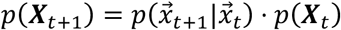, and due to the relationship 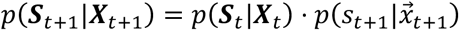, Eq. S3 can be written as follows:

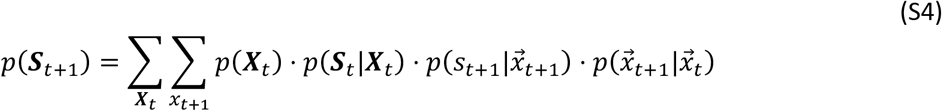

Using Bayes law and rearranging yields

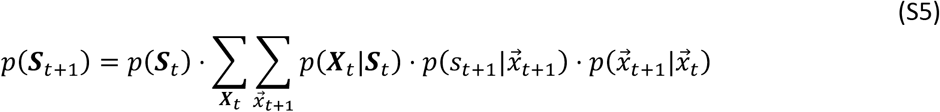

Re-writing the Markov decoder from the *Online Methods* (Eq. 2) using this notation,

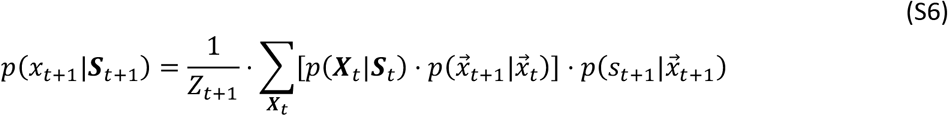

where the normalization factor *Z*_*t*+1_ satisfies the demand

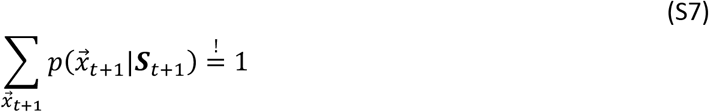

Summing over 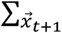 on both sides of Eq. S6 and rearranging we obtain

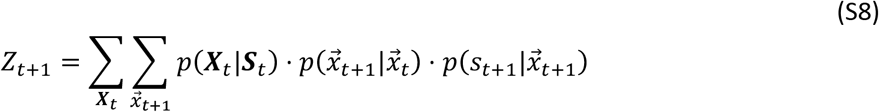

which is identical to the expression in Eq. S5. Thus, and in recursion,

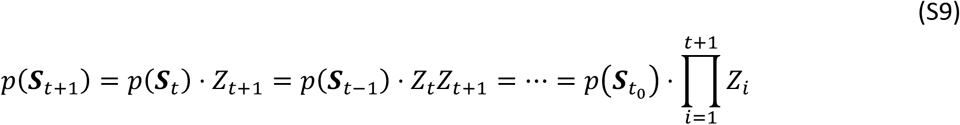

Finally, applying natural logarithm to Eq. S9 while neglecting the border term *p*(***S**_t_0__*), which is justified once sufficient iterations have been made, yields

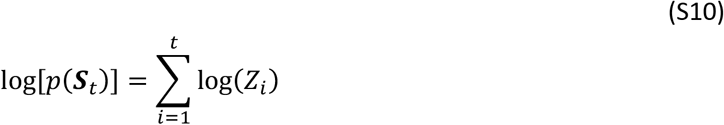

Qualitatively, the likelihood is accrued in a linear fashion over time, indicating that log(*Z*) was drawn from an approximately stationary distribution within each recording session (Supplementary Fig. 12). Therefore, to compare results between light and dark trials, we define the likelihood of simultaneous recorded spike trains as the mean of log(*Z*) over time, which can be thought as the amount of likelihood per time unit,

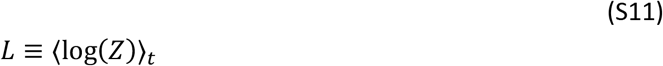

### b. Rate-adjusted likelihood

In this subsection we elucidate analytically why the posterior likelihood depends on the mean firing rate, and counter-intuitively decreases with increased spiking activity.

Denote by *ξ* a series of time binned spike trains which can take only {0,1} values in each time bin Δ*t*. Under the assumption of Poisson firing and in the limit of small Δ*t*, the posterior likelihood is written as

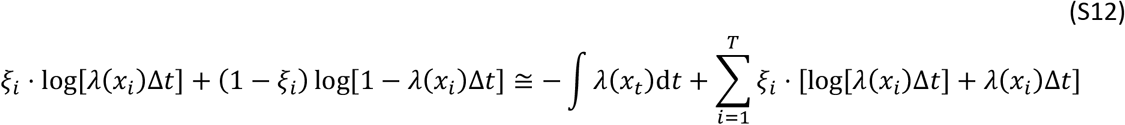

Since 0 < *λ*(*x_i_*)Δ*t* ≪ 1, ∀ *i*, we conclude (counter intuitively) that the addition of each spike can only decrease the posterior likelihood. This can be interpreted as arising from the fact that in the limit of small Δ*t*, the update of the likelihood with the addition of each spike is proportional to the probability that this particular spike will fall exactly within the relevant time bin.

Thus, to faithfully compare likelihoods between light and dark trials, it was essential to take into account mean firing rates modifications. The underlying assumption in the likelihood approach is that emitted spikes in darkness were generated from the same tuning curves as observed in light up to a multiplicative scaling factor. We down-sampled spike trains by randomly omitting spikes until the mean firing rates matched between the two trials, thus producing rate-adjusted spike trains. To demonstrate that the likelihood indeed depends on the mean firing rate and to justify that such random down sampling is appropriate, we varied a multiplicative factor which controlled the firing rate which accounts for the generation of simulated Poisson spike trains. As expected, the evaluated likelihood decreased as the mean firing rate increased but was completely unaffected if spike trains were subsequently down-sampled randomly using the corresponding multiplicative factor (Supplementary Fig. 13).

